# Parallel CRISPR-Cas9 screens reveal mechanisms of PLIN2 and lipid droplet regulation

**DOI:** 10.1101/2022.08.27.505556

**Authors:** Melissa A. Roberts, Kirandeep K. Deol, Mike Lange, Dara E. Leto, Alyssa J. Mathiowetz, Julian Stevenson, Sayed Hadi Hashemi, David W. Morgens, Emilee Easter, Kartoosh Heydari, Mike A. Nalls, Michael C. Bassik, Martin Kampmann, Ron R. Kopito, Faraz Faghri, James A. Olzmann

## Abstract

Lipid droplets (LDs) are lipid storage organelles that consist of a central core of neutral lipids surrounded by a phospholipid monolayer decorated with a unique set of integral and peripheral proteins. Invariably, at least one member of the perilipin family of proteins (PLIN1-5) associates with LDs in all cell types. Despite key roles of PLIN2 in governing hepatic lipid metabolism, the mechanisms that regulate PLIN2 levels remain incompletely understood. Here, we develop a set of genome-edited PLIN2 reporter cell lines that facilitate the analysis of genes that regulate PLIN2 and LD abundance. Leveraging these reporter cells in a series of CRISPR-Cas9 loss-of-function screens, we generate a comprehensive inventory of genes that influence PLIN2 levels under different metabolic conditions. Moreover, we uncouple their effects on PLIN2 expression and post-translational stability. Identified genetic modifiers include canonical genes that control LD metabolism (e.g., *ACSL3*, *DGAT2*, *PNPLA2*, *ABHD5*) as well as genes with less characterized roles in PLIN2 and LD regulation such as ubiquitination machinery (e.g., *MARCH6*, *UBE2J2*), transcription regulators (e.g., *HNF4A*, *HDAC3*), mitochondrial pathways (e.g., electron transport chain and mitochondrial fatty acid synthesis), and others. These CRISPR screens, and several published screens that focus on different aspects of lipid metabolism, provide the foundation for CRISPRlipid (http://crisprlipid.org), a versatile, online data commons for lipid-related functional genomics data. Together, our study uncovers new mechanisms of PLIN2 regulation and provides an extensive, phenotype-rich resource for the exploration of LD biology and lipid metabolism.

## INTRODUCTION

Lipid droplets (LDs) are the primary cellular organelle for lipid storage (Olzmann and Carvalho, 2019; Walther et al., 2017). LDs form at the endoplasmic reticulum (ER) through a stepwise process involving neutral lipid synthesis, phase separation of neutral lipids to form a lens structure within the ER bilayer, and directional emergence or budding of the LD from the cytosolic leaflet of the ER membrane (Renne et al., 2020; Thiam and Ikonen, 2021). These processes are catalyzed by ER proteins such as seipin, which forms an oligomeric complex that mediates triacylglycerol (TAG) aggregation to nucleate sites of LD biogenesis (Renne et al., 2020; Thiam and Ikonen, 2021). As hubs of lipid metabolism, LDs provide a dynamic lipid repository that can safely sequester lipids to prevent lipotoxicity and can be mobilized upon cellular demand to provide substrates for the generation of energy (i.e., β-oxidation) or the biosynthesis of lipid signaling molecules and membranes (Olzmann and Carvalho, 2019; Walther et al., 2017).

LDs consist of a core of neutral lipids, mostly triacylglycerol (TAG) and cholesteryl esters, surrounded by a phospholipid monolayer. Proteins are absent from the core of LDs, and a distinctive set of LD integral and peripheral proteins (i.e., the LD proteome) associates with the surrounding LD phospholipid monolayer and regulates LD functions, including biogenesis, breakdown, and interactions with neighboring organelles (Bersuker and Olzmann, 2017; Krahmer et al., 2013; Liu et al., 2004; Roberts and Olzmann, 2020). The perilipin proteins (PLIN1-5) are abundant LD proteins that are often referred to as LD coat proteins, and at least one PLIN family member is invariably found on LDs in all cell types (Kimmel and Sztalryd, 2016; Najt et al., 2022). The PLIN proteins share sequence homology, including a series of 11-mer repeats that mediate LD insertion and anchoring (Kimmel and Sztalryd, 2016; Najt et al., 2022). PLIN2, 3, and 5 also contain a C-terminal four-helix bundle and a unique α–β domain that may bind lipids (Kimmel and Sztalryd, 2016; Najt et al., 2022). PLIN proteins exhibit distinct tissue distributions, and while most regulate lipolysis, the different family members have additional unique functions that include mediating LD-organelle interactions at membrane contact sites, integrating nutrient signaling cascades that the regulate LD dynamics, and regulating the generation of lipid signaling molecules (Kimmel and Sztalryd, 2016; Najt et al., 2022).

PLIN2 is a ubiquitously expressed protein that plays important roles in regulating LDs, particularly in non-adipose tissues such as the liver. Global PLIN2 knockout (KO) mice exhibit reduced liver TAG levels and steatosis as well as resistance to diet-induced obesity (Orlicky et al., 2019). Moreover, liver-specific deletion of PLIN2 reduces liver steatosis, fibrosis, and inflammation, potentially by enhancing lipoprotein secretion and fatty acid oxidation while simultaneously suppressing *de novo* lipogenesis (Griffin et al., 2021; Najt et al., 2016). These beneficial phenotypes may also be explained by PLIN2’s competition with the rate-limiting enzyme in lipolysis, PNPLA2 (also known as ATGL), for LD association, which consequently inhibits lipolysis (Listenberger et al., 2007). In humans, an S251P polymorphism in PLIN2 has been connected to non-alcoholic steatohepatitis (NASH) and high serum lipid levels, but also with insulin sensitivity (Faulkner et al., 2020; Magné et al., 2013; Sentinelli et al., 2016). In addition, overexpression of PLIN2 increases LDs in a wide variety of tissues and high levels of PLIN2 are associated with several diseases, including hepatic steatosis (Gluchowski et al., 2017; Straub et al., 2008), conditions associated with high amounts of inflammation (Marschallinger et al., 2020), and several forms of cancer (Cruz et al., 2020). Thus, PLIN2 and LDs exhibit a reciprocal relationship, with LDs stabilizing PLIN2 and PLIN2 stabilizing LDs (Xu et al., 2019).

The influence of PLIN2 levels on LD metabolism and the connections of PLIN2 with disease pathogenesis underscore the importance of achieving a comprehensive understanding of the mechanisms that govern PLIN2 levels. Here, we performed a series of parallel CRISPR- Cas9 loss-of-function screens in a set of genome-edited PLIN2 reporter cells. These screens provide a phenotype-rich resource detailing the genetic modifiers that regulate PLIN2 and LDs, including regulators of PLIN2 expression and protein stability. Finally, using data from our screens related to PLIN2 regulation together with published lipid-focused screens, we establish CRISPRlipid (http://crisprlipid.org/) as an extensible, community-driven data commons for the exploration and comparison of data from functional genomics screens related to lipid biology.

## RESULTS

### Generation of a genome-edited PLIN2-GFP reporter cell line

To facilitate the analysis of PLIN2 regulation, CRISPR-Cas9 genome editing was used to generate a fluorescence-based PLIN2 reporter cell line in Huh7 hepatocellular carcinoma cells (**Figure 1A**). In this cell line, PLIN2 is endogenously tagged with enhanced green fluorescent protein (GFP) and an S-tag at its C-terminus (PLIN2-GFP) (**Figure 1A**). Endogenous tagging is essential to retain the normal regulation and levels of PLIN2 expression and to avoid artifacts associated with PLIN2 overexpression, such as the aberrant stabilization of LDs due to PLIN2 competition with PNPLA2 for LD binding (Kaushik and Cuervo, 2015; Listenberger et al., 2007). The edited PLIN2-GFP cells were distinguishable from parental cells by their increased GFP fluorescence using flow cytometry (**Figure 1B**). Immunoblotting for PLIN2 revealed the presence of both endogenous, untagged PLIN2 and GFP-tagged PLIN2 (**Figure 1C**, *middle blot*), and immunoblotting for GFP confirmed the presence of PLIN2-GFP as well as the absence of any free GFP (**Figure 1C**, *top blot*). These data indicate a heterozygous insertion of GFP in frame with PLIN2. Importantly, PLIN2-GFP properly localized to the LD membrane encircling the neutral lipid core of LDs (**Figure 1D**).

**Figure 1.**
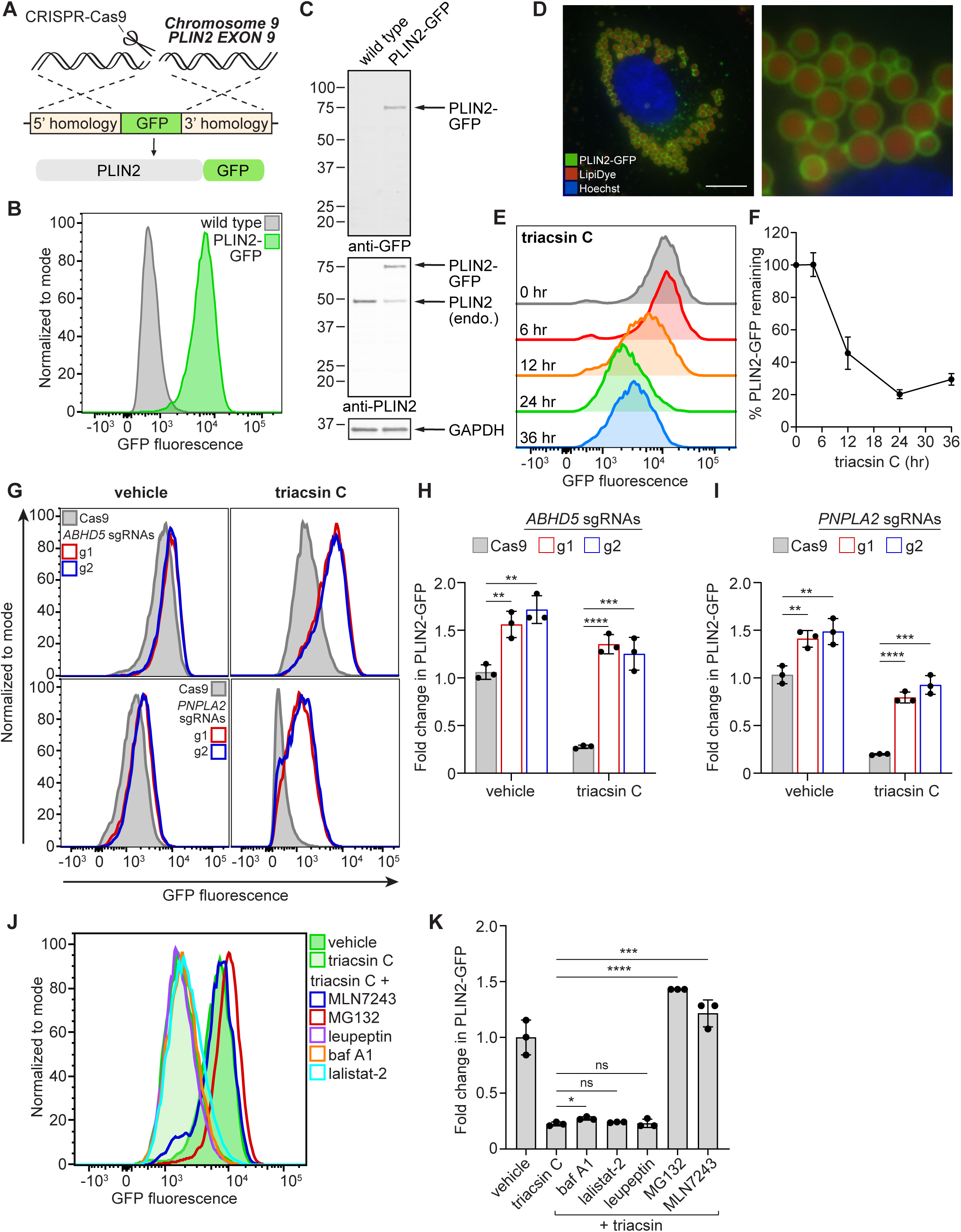
Characterization of a genetically-engineered PLIN2-GFP reporter cell line. **(A)** Illustration of CRISPR-Cas9 strategy used to generate the PLIN2-GFP knock-in reporter cell line. **(B)** Flow cytometry histogram of Huh7 wild type and PLIN2-GFP knock-in cells. **(C)** Immunoblot of parental and PLIN2-GFP Huh7 cells. Cells were lysed in SDS and proteins separated by SDS-PAGE and immunoblotted with antibodies against GFP, PLIN2, and GAPDH. **(D)** Representative fluorescence microscopy image of Huh7 PLIN2-GFP cells treated with 200 µM oleate for 24 hr. Lipid droplets were stained with 500 nM Lipi-Deep Red and nuclei with 5 µg/mL Hoechst 33342. Scale bar represents 10 µm. **(E)** Representative flow cytometry histograms of PLIN2-GFP cells treated with 1 µg/ml triacsin C for the indicated times. **(F)** Quantification of mode GFP fluorescence intensity from (E). Data represent mean ± SD of three biological replicates. **(G)** Representative flow cytometry histograms of PLIN2-GFP Cas9 expressing cells with no sgRNAs (grey) or sgRNAs against *ABHD5* (top, red and blue) or *PNPLA2* (bottom, red and blue). Cells were treated with 1 µg/ml triacsin C or DMSO for 24 hr. **(H)** Quantification of the fold change in mode GFP fluorescence intensity in PLIN2-GFP Cas9 cells versus cells expressing sgRNAs against *ABHD5* from (G). Data represent mean ± SD of three biological replicates. ** p < 0.01, *** p <0.001, **** p <0.0001 by two-tailed, unpaired t-test. **(I)** Quantification of the fold change in mode GFP fluorescence intensity in PLIN2-GFP Cas9 cells versus cells expressing sgRNAs against *PNPLA2* from (G). Data represent mean ± SD of three biological replicates. ** p < 0.01, *** p <0.001, **** p <0.0001 by two-tailed, unpaired t-test. **(J)** Representative flow cytometry histogram of PLIN2-GFP cells treated with 1 µg/ml triacsin C or 1 µg/ml triacsin C with 10 µM MLN7243, 10 µM MG132, 50 µM leupeptin, 250 nM bafilomycin A1, 10 µM lalistat-2, or DMSO for 24 hr. **(K)** Quantification of fold change in mode GFP fluorescence intensity from (J). Data represent mean ± SD of three biological replicates. * p < 0.05, *** p = 0.0001, **** p < 0.0001 by two-tailed, unpaired t-test.

To further characterize the PLIN2-GFP reporter cell line, we examined the effects of oleate-stimulated LD biogenesis and triacsin C treatment. Triacsin C is an acyl-CoA synthetase inhibitor that blocks TAG biosynthesis and starves cells of activated fatty acids, triggering LD breakdown. Consistent with the strong correlation between PLIN2 levels and LD abundance, endogenous PLIN2 and PLIN2-GFP levels increased following oleate treatment and decreased in response to triacsin C treatment (**Figure 1E,F; Figure S1A,B**). As expected, the changes in PLIN2-GFP levels correlated with the changes in the fluorescence of the neutral lipid stain monodansylpentane (MDH) and side scatter (**Figure S1B**). To define the mechanisms that regulate PLIN2-GFP decrease during lipolysis, we introduced single guide RNAs (sgRNAs) targeting the TAG lipase *PNPLA2* (also known as ATGL) and its co-activator *ABHD5* (also known as CGI-58) into PLIN2-GFP cells expressing Cas9. Depletion of PNPLA2 or ABHD5 led to a small increase in PLIN2-GFP levels under basal conditions and a dramatic block in the PLIN2-GFP decrease in response to triacsin C treatment (**Figure 1G-I**), indicating that LD degradation during triacsin C treatment requires PNPLA2 and ABHD5-dependent lipolysis. These findings highlight the differential impact of canonical LD regulatory genes depending on the metabolic state of the cell, underscoring the importance of characterizing LD regulation under different metabolic conditions.

PLIN2 is stabilized by its insertion into LDs and uninserted PLIN2 is degraded by the ubiquitin-proteasome system (UPS) in cell types that have few LDs, such as HEK293 and HeLa (Masuda et al., 2006; Nguyen et al., 2019; Xu et al., 2005). In contrast to HEK293 and HeLa cells, Huh7 cells have large amounts of PLIN2-positive LDs. Whether this pool of LD-inserted PLIN2 in Huh7 cells is degraded by the proteasome or an alternative pathway, such as via autophagy or the lysosome, during the lipolytic consumption of LDs is unknown. During triacsin C-induced LD degradation, the decrease in PLIN2-GFP levels was strongly blocked by co-incubation with MG132, an inhibitor of the proteasome, or MLN7243, an inhibitor of the E1 ubiquitin-activating enzyme (**Figure 1J,K; Figure S1C,D**). In contrast, co-incubation with leupeptin, bafilomycin A1 (baf A1), or lalistat-2, inhibitors of lysosomal functions, had no effect (**Figure 1J,K**). These findings indicate that the clearance of LD-inserted PLIN2 during lipolysis requires the UPS and not lysosomal degradation pathways such as lipophagy and chaperone-mediated autophagy (CMA). Together, these data demonstrate that the genome-edited PLIN2-GFP Huh7 cell line provides a reliable reporter of PLIN2 levels and LD dynamics.

### CRISPR-Cas9 screens identify metabolic state-dependent regulators of PLIN2-GFP

To systematically profile the genetic factors that regulate PLIN2 levels, we performed genome-wide CRISPR-Cas9 loss-of-function screens under basal and lipolytic (i.e., triacsin C treated) conditions (**Figure 2A**). Cas9 expressing PLIN2-GFP reporter cells were infected with a lentiviral sgRNA library containing 10 sgRNAs per gene (Morgens et al., 2016, 2017) and the top and bottom 30 percent of PLIN2-GFP fluorescent cells were isolated by FACS. For the lipolysis screen, cells were pretreated with triacsin C for 24 hr prior to FACS. The high and low GFP fluorescent cells represent cells in which gene disruption increased or decreased PLIN2-GFP levels, respectively. Following deep sequencing of PCR-amplified library barcodes, sgRNA enrichment was analyzed using Cas9 high-Throughput maximum Likelihood Estimator (casTLE) (Morgens et al., 2016, 2017). Employing a 10 percent false discovery rate (FDR), we identified 192 significant gene hits in the basal screen and 275 significant gene hits in the lipolytic screen (**Table S1, Figure S2A,B**). *PLIN2* itself was identified as a hit since disruption of the endogenous gene reduces PLIN2-GFP expression (**Figure S2A,B**) and many canonical LD regulators were identified as high-confidence hit genes with strong effects on the amount of PLIN2-GFP (**Figure S2A,B**), consistent with the reciprocal stabilization of PLIN2 and LDs. For example, genes required for TAG breakdown, such as *PNPLA2* and *ABHD5*, were identified as hits that increased PLIN2-GFP in the basal and lipolytic screens (**Figure S2A,B**), and genes required for TAG synthesis, such as *DGAT2* and *ACSL3*, were identified as hits that decreased PLIN2-GFP in the basal screen (**Figure S2A,B**).

**Figure 2.**
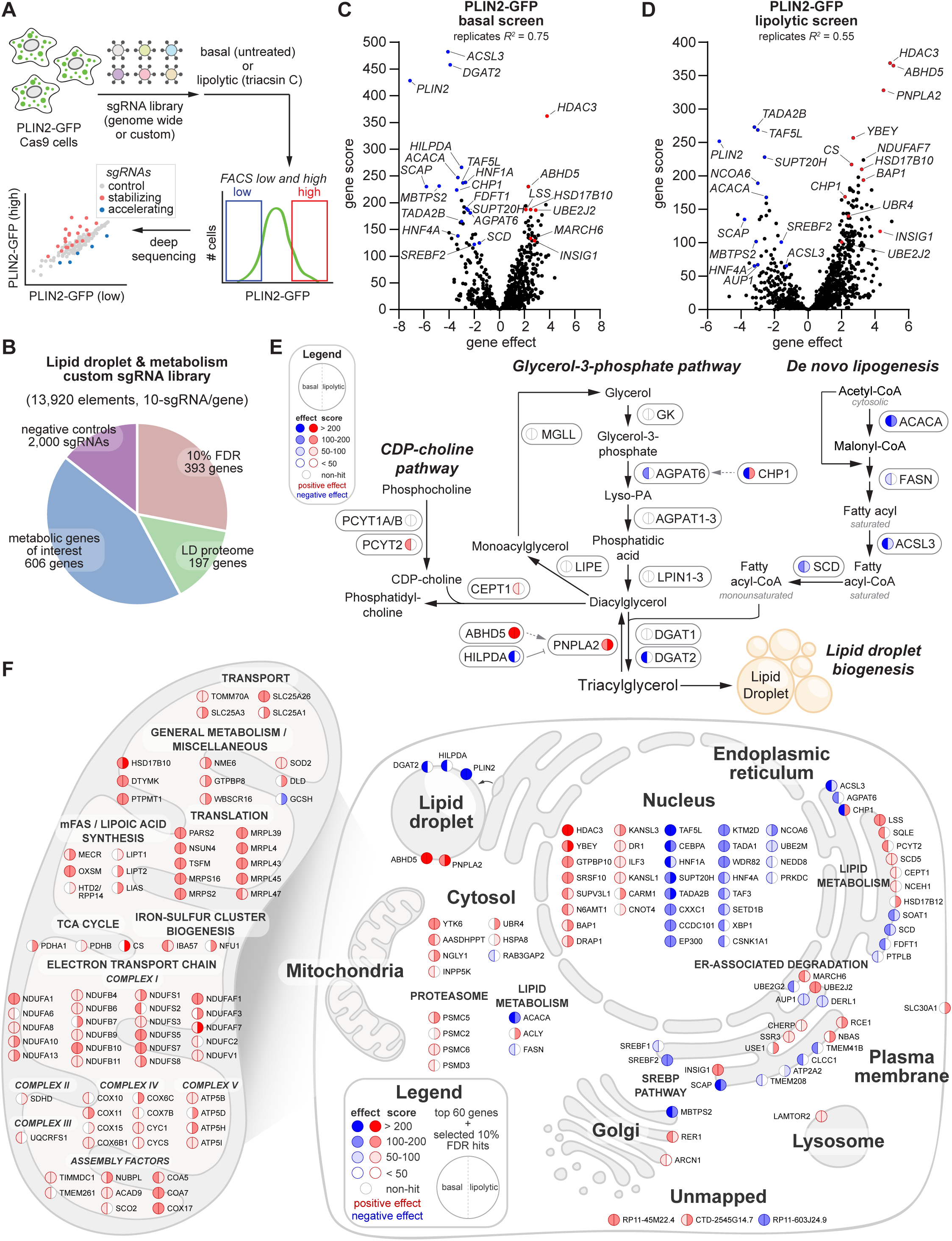
CRISPR-Cas9 screen identifies metabolic state-dependent regulators of PLIN2-GFP. **(A)** Schematic of FACS-based CRISPR-Cas9 screening workflow. **(B)** Breakdown of the custom 10 guide-per-gene Lipid Droplet and Metabolism sgRNA library generated for batch retest screens. **(C,D)** Volcano plots indicating the gene effects and gene scores for individual genes from batch retest CRISPR-Cas9 screens of PLIN2-GFP cells under basal (C) and triacsin C treated (D) conditions. Gene effects and gene scores were computed from two biological replicates per screen. **(E)** Schematic of the triacylglycerol synthesis and breakdown pathways. Genes are annotated with nodes corresponding to gene effects and scores from batch retest basal and triacsin C screens. **(F)** Cell map showing putative regulators of PLIN2 and LDs under basal and lipolytic (triacsin C treated) conditions. Map consists of the top 60 genes (ranked by gene score) from each screen and selected genes that met a 10 percent false discovery rate cutoff. Cellular localizations and functional groupings were assigned based on Gene Ontology annotations and/or previous literature.

To characterize and validate our hits, we performed a set of duplicate high coverage batch retest screens, which are known to reduce false positives and false negatives (Bassik et al., 2013; Dubreuil et al., 2020; Han et al., 2017; Parnas et al., 2015). For these screens, we synthesized a custom sgRNA library (10 sgRNAs per gene) targeting genes passing a 10% FDR from our genome-wide CRISPR screens, genes encoding proteins present in prior LD proteomic analyses (Bersuker et al., 2018), and selected genes of interest with roles related to lipid metabolism (**Figure 2B**). In total, this new “Lipid Droplet and Metabolism” sgRNA library contains 13,920 sgRNAs with 11,920 gene targeting sgRNAs and 2,000 negative control sgRNAs (**Figure 2B; Table S2**). PLIN2-GFP cells were infected with the custom sgRNA library and duplicate high coverage screens performed under basal and lipolytic conditions, as described for the genome-wide screen (**Figure 2A**). The batch retest screens resulted in higher gene effects (a score of the phenotype strength) and gene scores (a confidence metric) relative to the genome-wide screens (**Figure 2C,D**; **Figure S2A,B**), strongly correlated with the genome-wide screens (**Figure S2C,D**), identified expected LD regulatory genes (**Figure 2C,D**; **Figure S2E**), and further highlighted differential gene effects under basal and lipolytic conditions (**Figure S2F**).

Enzymes involved in nearly every step of neutral lipid synthesis were identified (**Figure 2E**). The depletion of enzymes that mediate neutral lipid synthesis reduced PLIN2-GFP, including the *de novo* fatty acid synthesis enzymes *ACACA*, *ACLY*, and *FASN*, the fatty acid desaturase *SCD*, the acyl-CoA synthetase *ACSL3*, the glycerol-3-phosphate acyltransferase *AGPAT6* (also known as GPAT4) and its activator *CHP1*, and the diacylglycerol acyltransferase *DGAT2* (**Figure 2E**). Conversely, disruptions in genes involved in neutral lipid degradation increased PLIN2-GFP, including *PNPLA2* and *ABHD5*, while the disruption of the PNPLA2 inhibitor *HILPDA* reduced PLIN2-GFP (**Figure 2E**).

A cellular map of the top gene hits from the batch retest screens provides a visual representation of selected high-confidence PLIN2 regulators, highlighting expected regulators of LD biology such as neutral lipid synthesis enzymes as well as numerous genes with no prior connection to LD metabolism or PLIN2 regulation (**Figure 2F**). Several genes that comprise the SREBP pathway were identified, including *SREBF1/2*, *SCAP*, *INSIG1, and MBTPS2* (**Figure 2F**). We validated these results for three of the genes, finding that targeted disruption of *INSIG1* increased PLIN2-GFP and disruption of *MBTPS2* or *SCAP* reduced PLIN2-GFP (**Figure S3A**). PLIN2 may be an SREBP target, or it could be post-translationally affected by SREBP-mediated alterations in cellular lipid metabolism and LD abundance. Although Huh7 LDs primarily contain TAG, multiple genes involved in cholesterol metabolism were also observed as PLIN2 regulators (e.g., *LSS*, *SQLE*, *SOAT1*, *FDFT1*). These cholesterol metabolic genes were distributed across the cholesterol synthesis pathway (pathway map in **Figure S3B**) and may influence PLIN2 abundance by regulating SREBP signaling or by contributing cholesteryl esters that are sequestered in LDs. We detected a remarkable number of genes involved in mitochondrial biology (**Figure 2F**). In all cases the disruption of these genes increased PLIN2-GFP (**Figure 2F**). Particularly striking was the enrichment of genes within the electron transport chain (**Figure 2F**). One possibility is that disruptions in the electron transport chain reduce fatty acid oxidation and LD breakdown. An alternative is that the increase in LDs reflects a response to the induction of mitochondrial stress pathways, as LD biogenesis is known to increase in response to diverse cellular stresses. Interestingly, many enzymes involved in the mitochondrial fatty acid synthesis (mFAS) and lipoic acid synthesis pathways were identified (**Figure 2F,** pathway map in **Figure S3C**). While disruption in the cytosolic fatty acid synthesis pathway reduced PLIN2-GFP, likely by reducing the production of *de novo* fatty acids required for neutral lipid synthesis, the disruption of the mitochondrial fatty acid synthesis pathway increased PLIN2-GFP (**Figure 2F**; **Figure S3C**). The mitochondrial fatty synthesis pathway generates the eight-carbon saturated fatty acid acid caprylic acid (octanoic acid), which is converted into lipoic acid for the lipoylation of several important mitochondrial enzymes, such as pyruvate dehydrogenase. Indeed, several components of the pyruvate dehydrogenase complex are detected as high confidence hits – *PDHA1*, *PDHB*, *DLAT*, and *DLD* (**Figure 2F**; **Figure S3C**). The mitochondrial fatty synthesis pathway also generates longer acyl chains and has roles in regulating electron transport chain assembly and function, independent of protein lipoylation (Nowinski et al., 2020). While the role of the mitochondrial fatty synthesis pathway in controlling electron transport function may contribute to our observed PLIN2-GFP phenotypes, the identification of lipoyl synthase *LIAS* and both lipoyltransferases, *LIPT1* and *LIPT2,* suggest that mitochondrial enzyme lipoylation is important. Thus, our genome-wide and batch retest screens provide a comprehensive inventory of genetic modifiers that govern PLIN2 levels under basal and lipolytic metabolic states.

### MARCH6 regulation of neutral lipid storage in LDs indirectly stabilizes PLIN2

PLIN2 levels are controlled by its degradation via chaperone-mediated autophagy (Kaushik and Cuervo, 2015, 2016) and the ubiquitin-proteasome system (Masuda et al., 2006; Nguyen et al., 2019; Xu et al., 2005). In cell lines that have low amounts of LDs, uninserted PLIN2 is degraded through a mechanism involving the ER-resident E3 ubiquitin-protein ligase MARCH6 (also known as TEB4) (Nguyen et al., 2019). We detected several sgRNAs targeting *MARCH6* and its cognate E2 ubiquitin-conjugating enzyme *UBE2J2* under basal (**Figure 3A,D**) and lipolytic conditions (**Figure S4A,B**), identifying these genes as high confidence PLIN2 regulators and suggesting that MARCH6 and UBE2J2 cooperate to control PLIN2 stability in LD-rich Huh7 cells. Indeed, the disruption of *MARCH6* or *UBE2J2* with 3 or 2 independent sgRNAs, respectively, increased PLIN2-GFP levels by flow cytometry under basal conditions and more strongly under triacsin C treated conditions (**Figure 3B,C,E,F**). Due to the lack of effective anti-MARCH6 antibodies, MARCH6 depletion was confirmed by genome sequencing and the accumulation of its known substrate squalene monooxygenase (SQLE) (Foresti et al., 2013; Gill et al., 2011; Stevenson et al., 2016) (**Figure S4C**), and UBE2J2 knockout was directly confirmed by immunoblotting (**Figure S4D**).

**Figure 3.**
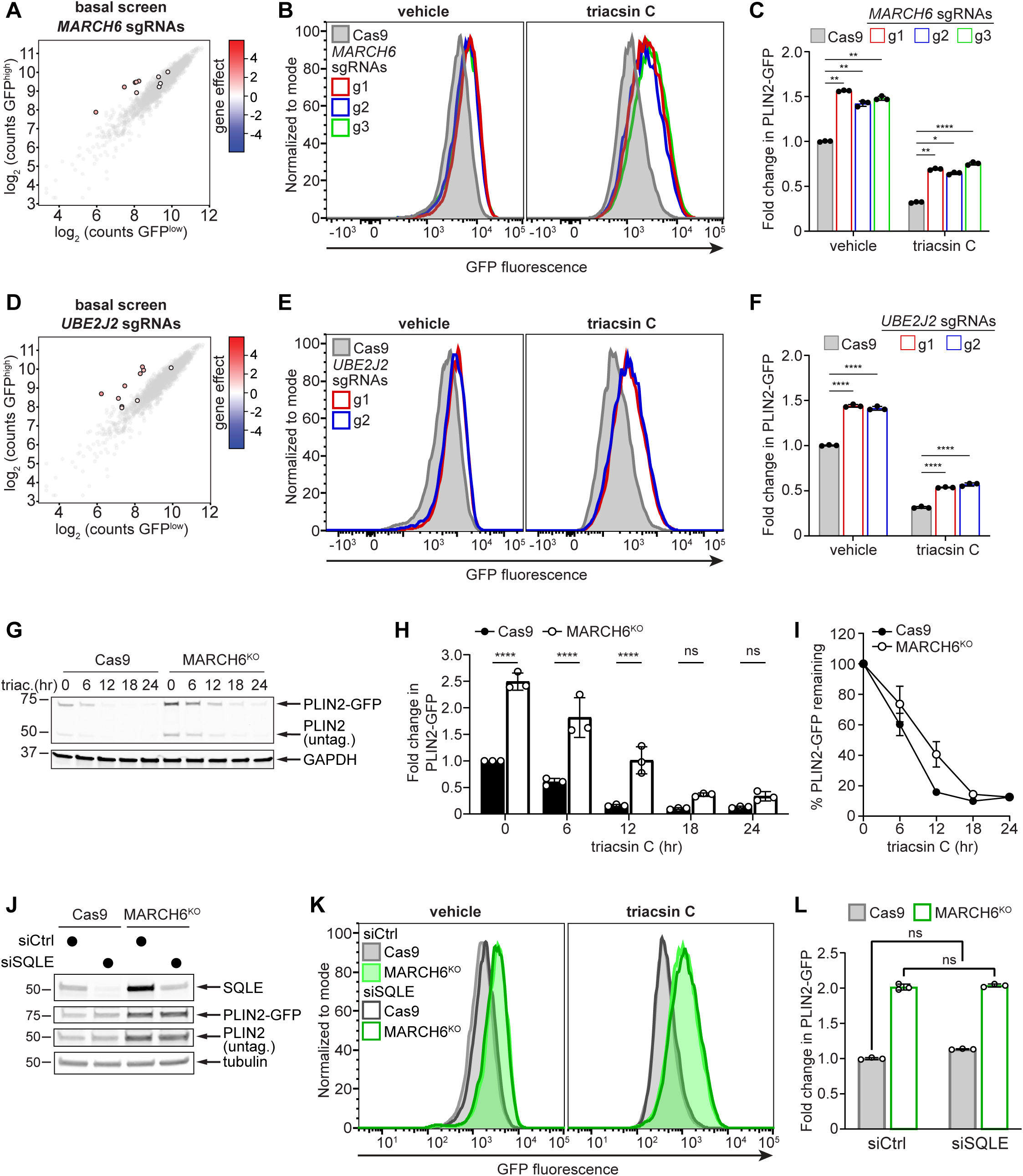
Regulation of PLIN2-GFP levels by the E3 ligase MARCH6. **(A)** Cloud plot indicating deep sequencing counts corresponding to *MARCH6* (color scale) or negative control sgRNAs (grey scale) from one replicate of the PLIN2-GFP basal batch retest screen. **(B)** Representative flow cytometry histograms of PLIN2-GFP Cas9 cells expressing no sgRNAs (grey) or three different sgRNAs against *MARCH6* (colors) following treatment with 1 µg/ml triacsin C or DMSO for 24 hr. **(C)** Quantification of the fold change in mean GFP fluorescence intensity from (B). Data represent mean ± SD of three biological replicates. * p < 0.05, ** p < 0.01, **** p < 0.0001 by two-tailed, unpaired t-test. **(D)** Cloud plot indicating deep sequencing counts corresponding to *UBE2J2* (color scale) or negative control sgRNAs (grey scale) from one replicate of the PLIN2-GFP basal batch retest screen. **(E)** Representative flow cytometry histograms of PLIN2-GFP Cas9 cells expressing no sgRNAs (grey) or two different sgRNAs against *UBE2J2* (colors) following treatment with 1 µg/ml triacsin C or DMSO for 24 hr. **(F)** Quantification of the fold change in mean GFP fluorescence intensity from (E). Data represent mean ± SD of three biological replicates. **** p < 0.0001 by two-tailed, unpaired t-test. **(G)** Representative immunoblot of PLIN2-GFP Cas9 and PLIN2-GFP MARCH6^KO^ cells treated with 1 µg/ml triacsin C for the indicated times. Cells were lysed in SDS and proteins separated by SDS-PAGE and immunoblotted with antibodies against PLIN2 and GAPDH. **(H)** Quantification of the fold change in PLIN2-GFP protein levels in Cas9 and MARCH6^KO^ cells at the indicated triacsin C treatment times from (G). Data represent mean ± SD of three biological replicates. **** p < 0.0001. **(I)** Quantification of PLIN2-GFP protein levels from (G). PLIN2-GFP protein levels were normalized to levels at time 0 hr for each cell line. Data represent mean ± SD of three biological replicates. **(J)** Immunoblot of PLIN2-GFP Cas9 or MARCH6^KO^ cells incubated with *SQLE* or control siRNAs for 72 hr. Cells were lysed in SDS and proteins separated by SDS-PAGE and immunoblotted with antibodies against SQLE, PLIN2, and tubulin. **(K)** Representative flow cytometry histograms of PLIN2-GFP Cas9 and PLIN2-GFP MARCH6^KO^ cells incubated with *SQLE* or control siRNAs for 72 hr and treated with 1 µg/ml triacsin C or DMSO for 24 hr. **(L)** Quantification of the fold change in mean GFP fluorescence intensity from (K). Data represent mean ± SD of three biological replicates. No significant changes were observed by two tailed, unpaired t-test.

To characterize the role of MARCH6 in regulating PLIN2 levels, we analyzed the turnover kinetics of PLIN2-GFP and endogenous, untagged PLIN2 in Cas9 expressing control and MARCH6 knockout (MARCH6^KO^) cells during a 24 hr triacsin C treatment. PLIN2-GFP and untagged PLIN2 exhibited higher steady state levels in MARCH6^KO^ cells (**Figure 3G,H**). Indeed, PLIN2-GFP levels were higher in MARCH6^KO^ cells relative to control cells at all time points throughout the triacsin C treatment time course (**Figure 3G,H**). However, quantification of the PLIN2-GFP turnover rate after normalization to the initial higher levels indicates that the rate of PLIN2-GFP degradation during lipolysis is unaltered in MARCH6^KO^ cells (**Figure 3I**). The endogenous untagged PLIN2 responded similarly to the GFP-tagged protein, showing increased levels in the MARCH6^KO^ cells (**Figure S4E**) and an unaltered rate of clearance (**Figure S4F**). We considered the possibility that the increase in SQLE, which is a rate-limiting enzyme in cholesterol biosynthesis (Foresti et al., 2013; Gill et al., 2011; Stevenson et al., 2016), could be responsible for the changes in PLIN2 levels in the MARCH6^KO^ cells. However, siRNA-mediated depletion of SQLE had no effect on the levels of untagged PLIN2 or PLIN2-GFP in western blots (**Figure 3J**) or of PLIN2-GFP levels measured by flow cytometry under basal or triacsin C treated conditions (**Figure 3K,L**). Together, these data indicate that although MARCH6 affects PLIN2 levels in Huh7 cells, MARCH6 is not required for the clearance of LD-inserted PLIN2 during the lipolytic breakdown of LDs and its effects on PLIN2 are independent of its known substrate SQLE.

Given the increase in PLIN2 levels in MARCH6^KO^ cells and the high correlation between PLIN2 and LDs, we examined LDs in MARCH6^KO^ cells by confocal imaging (**Figure 4A,B**). Depletion of MARCH6 in parental Huh7 cells increased endogenous PLIN2 levels (**Figure S5A**) and the amount of LDs under basal and triacsin C treated conditions (**Figure 4A,B, Figure S5B**). LD size also increased under basal conditions, but only one cell line exhibited a statistical difference in LD area under the triacsin C treated conditions (**Figure 4A,B, Figure S5B**). To gain insight into the biochemical changes in cellular lipids, we performed untargeted lipidomics on our MARCH6^KO^ cells. Principal component analysis of the lipidomics data showed a distinct cluster of MARCH6^KO^ cell lines that was well separated from the control cells (**Figure 4C**). Consistent with our imaging data showing an increase in neutral lipids stored in LDs, the lipidomics analyses revealed a general increase in TAG species (**Figure 4D**). These TAG species were diverse in their compositions of conjugated fatty acids, including their degree of unsaturation and number of carbons (i.e., fatty acid chain length) (**Figure 4E, Figure S5D**). There were also reductions in phosphatidylcholine (PC) as well as increases in phosphatidylinositol, indicating that the effects of MARCH6 on the cellular lipid landscape are not limited to TAG (**Figure 4D,E, Figure S5E**).

**Figure 4.**
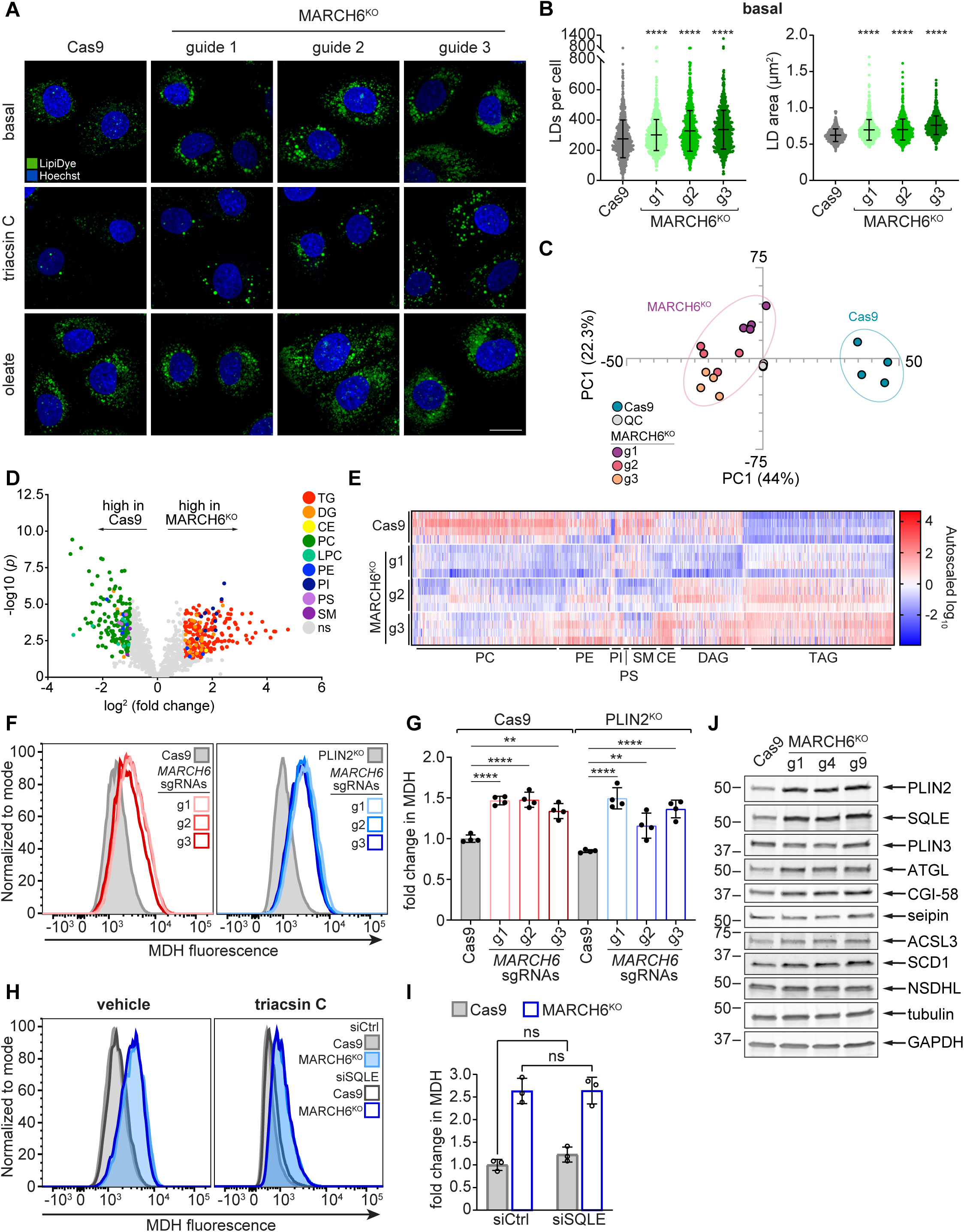
The E3 ligase MARCH6 regulates neutral lipid storage independently of PLIN2. **(A)** Representative confocal microscopy images of Cas9 control cells expressing sgRNAs against a non-coding region of the genome and three independent, clonally isolated MARCH6^KO^ cell lines. Cells were imaged at steady state (basal conditions) or following 8 hr of treatment with 1 µg/ml triacsin C or 200 µM oleate. Lipid droplets were stained with 500 nM Lipi-Green neutral lipid stain and nuclei with 5 µg/mL Hoechst 33342. Scale bar represents 20 µm. **(B)** Quantification of the number (left panel) and area (right panel) of basal lipid droplets per cell from (A). Data represent mean ± SD of three biological replicates. \*\*\*\**p* < 0.0001 by one-way ANOVA with Dunnett’s multiple comparisons test. **(C)** Principal component analysis plot (PCA) of untargeted lipidomics analysis of Cas9 control cells expressing sgRNAs against a non-coding region of the genome and three independent, clonally isolated MARCH6^KO^ cell lines. The analysis was performed on four biological replicates. **(D)** Volcano plot of untargeted lipidomics analysis of Cas9 control cells expressing sgRNAs against a non-coding region of the genome and three independent, clonally isolated MARCH6^KO^ cell lines. The analysis was performed on four biological replicates. TG = triacylglycerol; DG = diacylglycerol; CE = cholesterol ester; PC = phosphatidylcholine; LPC = lysophosphatidylcholine; PE = phosphatidylethanolamine; PI = phosphatidylinositol; PS = phosphatidylserine; SM = sphingomyelin; ns = non-significant. **(E)** Heat map indicating changes in lipid species from the untargeted lipidomics analysis of Cas9 control cells expressing sgRNAs against a non-coding region of the genome and three independent, clonally isolated MARCH6^KO^ cell lines. Data are autoscaled and normalized to the average abundance of each species presented. **(F)** Representative flow cytometry histograms of Cas9 control cells expressing sgRNAs against a non-coding region of the genome or three independent MARCH6^KO^ cell lines (left panel) or PLIN2^KO^ cells expressing sgRNAs against a non-coding region of the genome or three independent, clonally isolated MARCH6^KO^ cell lines (right panel). Neutral lipids were stained with 100 µM MDH and analyzed using the Pacific Blue channel. **(G)** Quantification of the fold change in mean MDH fluorescence intensity from (F). Data represent mean ± SD of three biological replicates. ** *p* < 0.01, **** *p* < 0.0001. **(H)** Representative flow cytometry histograms of PLIN2-GFP Cas9 control and PLIN2-GFP MARCH6^KO^ cells incubated with *SQLE* or control siRNAs for 72 hr and treated with 1 µg/ml triacsin C or DMSO for 24 hr. Neutral lipids were stained with 100 µM MDH and analyzed using the Pacific Blue channel. **(I)** Quantification of the fold change in mean MDH fluorescence intensity from (H). Data represent mean ± SD of three biological replicates. **(J)** Immunoblots of Huh7 Cas9 cells expressing sgRNAs against a non-coding region of the genome or three independent sgRNAs against *MARCH6* (pooled knockouts). Cells were lysed in SDS and proteins separated by SDS-PAGE and immunoblotted for the indicated proteins.

To test whether the higher amount of PLIN2 in the MARCH6^KO^ cells is responsible for the higher neutral lipid levels, we introduced MARCH6 sgRNAs into PLIN2^KO^ cell lines and measured neutral lipid content by flow cytometry. As anticipated, loss of PLIN2 reduced neutral lipid content (**Figure S5F-I**). However, the loss of PLIN2 had no effect on the increase in neutral lipid content following MARCH6 depletion (**Figure 4F,G, Figure S5J-L**). In addition, depletion of SQLE also had no effect on neutral lipid levels in the MARCH6^KO^ cell line (**Figure 4H,I**). These findings indicate that the effect of MARCH6 on neutral lipid levels is independent of PLIN2 and SQLE stabilization. These data support a model in which MARCH6 regulates TAG and LD levels, which in turn affect PLIN2 stability. Indeed, other obligate LD proteins that are also reciprocally regulated by LD levels are increased in the MARCH6^KO^ cells, including ATGL and CGI-58 (**Figure 4J**).

### Parallel screens uncouple transcriptional and post-translational mechanisms of PLIN2 regulation

The screens employing our PLIN2-GFP reporter cell line identified regulators that influence PLIN2 levels through diverse mechanisms – transcriptional, translational, and post-translational. To systematically uncouple these mechanisms of PLIN2 regulation, we engineered a new reporter cell line in which GFP and blue fluorescent protein (BFP), separated by a short P2A sequence, were integrated in frame with the C-terminus of endogenous PLIN2 (**Figure 5A**). The short self-cleaving P2A peptide sequence induces a ribosomal skip and allows BFP to be translated from the same transcript as the PLIN2-GFP fusion protein (**Figure 5A**). Thus, the BFP levels report on transcription and the GFP:BFP ratio reports on post-translational regulation.

**Figure 5.**
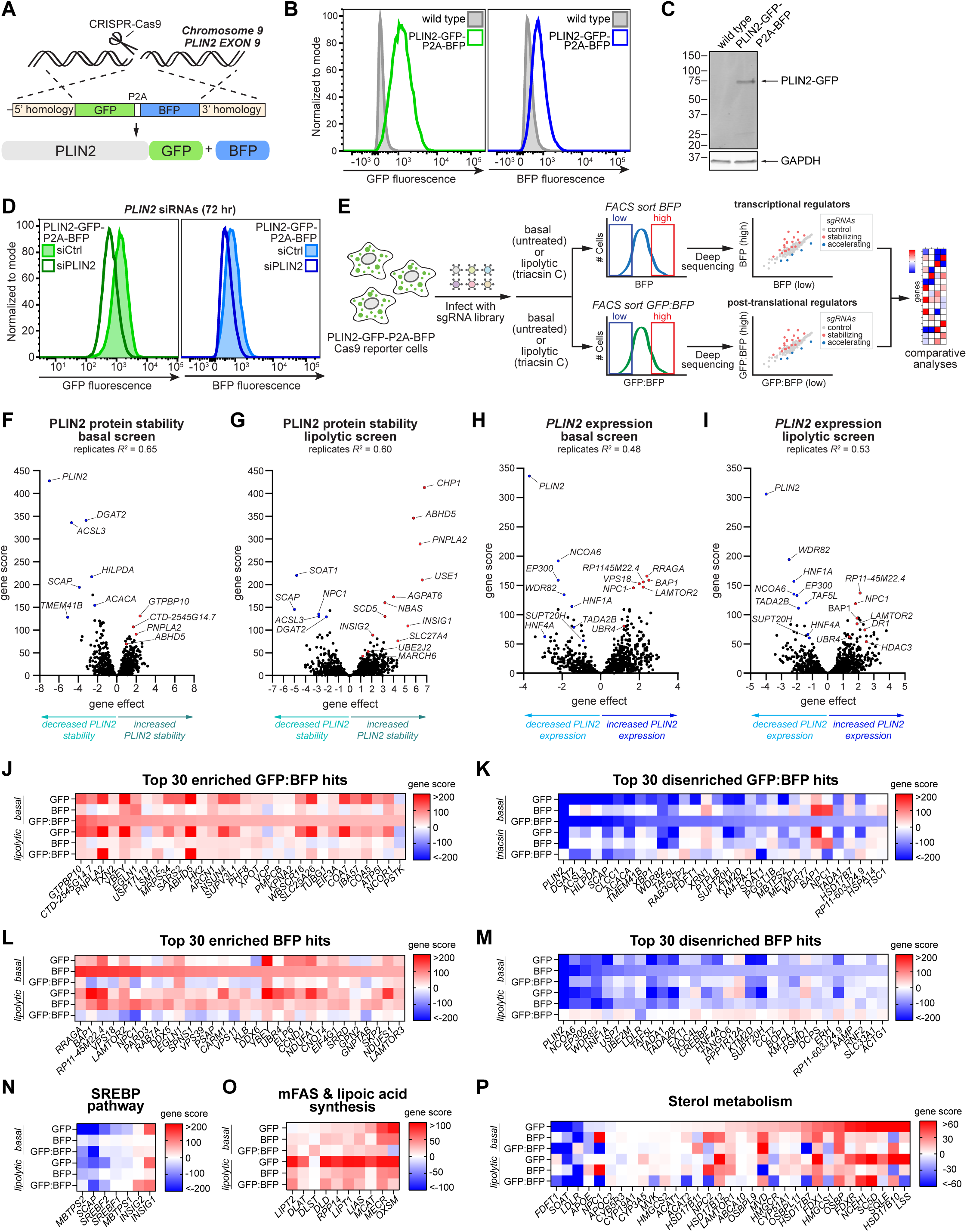
Analysis of pre- and post-translational PLIN2-GFP regulators. **(A)** Schematic of the CRISPR-Cas9 strategy used to generate the PLIN2-GFP-P2A-BFP reporter cell line. **(B)** Flow cytometry histograms of Huh7 parental and PLIN2-GFP-P2A-BFP knock-in cells. GFP and BFP were detected using FITC and Pacific Blue channels, respectively. **(C)** Immunoblot of Huh7 parental and PLIN2-GFP-P2A-BFP cells. Cells were lysed in SDS and proteins separated by SDS-PAGE and immunoblotted with antibodies against GFP and GAPDH. **(D)** Representative flow cytometry histograms of PLIN2-GFP-P2A-BFP cells incubated with *PLIN2* or control siRNAs for 72 hr. GFP and BFP were detected using FITC and Pacific Blue channels, respectively. **(E)** Schematic of batch retest CRISPR-Cas9 screen of PLIN2-GFP-P2A-BFP cells. **(F,G)** Volcano plots indicating the gene effects and gene scores for individual genes from GFP:BFP screens under basal (F) and triacsin C treated (G) conditions. Screens were performed in duplicate and volcano plots indicate combined gene effects and scores. **(H,I)** Volcano plots indicating the gene effects and gene scores for individual genes from BFP screens under basal (H) or triacsin C treated (I) conditions. Screens were performed in duplicate and volcano plots indicate combined gene effects and scores. **(J,K)** Heat maps displaying the signed gene scores of the top 30 enriched (J) and disenriched (K) genes (GFP:BFP^high^ relative to GFP:BFP^low^) from the GFP:BFP batch retest screen under basal conditions. Signed gene scores from all other batch retest screens are included for comparison. **(L,M)** Heat maps displaying the signed gene scores of the top 30 enriched (L) and disenriched (M) genes (BFP^high^ relative to BFP^low^) from the BFP batch retest screen under basal conditions. Signed gene scores from all other batch retest screens are included for comparison. **(N)** Heat map displaying the signed gene scores of SREBP pathway-related genes from batch retest GFP, BFP, and GFP:BFP screens. **(O)** Heat map displaying the signed gene scores of mitochondrial fatty acid synthesis (mFAS) and lipoic acid genes from batch retest GFP, BFP, and GFP:BFP screens. **(P)** Heat map displaying the signed gene scores of sterol metabolism-related genes from batch retest GFP, BFP, and GFP:BFP screens.

The PLIN2-GFP-P2A-BFP reporter cell line exhibited GFP and BFP fluorescence above wild type parental cells (**Figure 5B**) and a single PLIN2-GFP band was detected by immunoblotting at the expected molecular weight (**Figure 5C**), indicating in-frame insertion of GFP and appropriate cleavage of PLIN2-GFP and BFP at the P2A peptide. siRNA depletion of the PLIN2 transcript led to a reduction in both GFP and BFP (**Figure 5D**), consistent with BFP expression from the PLIN2 transcript. Oleate-induced LD biogenesis increased and triacsin C-induced lipolysis decreased PLIN2-GFP levels, but not BFP levels (**Figure S6A**), in agreement with the post-translational regulation of PLIN2-GFP stability by LDs. These data demonstrate the utility of this new reporter cell line in distinguishing between transcriptional and post-translational mechanisms of PLIN2 regulation.

Employing the PLIN2-GFP-P2A-BFP reporter cell line, we performed parallel high coverage screens using our Lipid Droplet and Metabolism sgRNA library (**Figure 5E**). Cells were sorted based upon BFP fluorescence to identify transcriptional regulators and the ratio of GFP:BFP fluorescence to identify post-translational regulators (**Figure 5E, Table S1**). As expected, sgRNAs targeting PLIN2 were disenriched in both screens (**Figure S6B,C**). Genes that are directly involved in the biosynthesis and turnover of neutral lipids and LDs were identified in the GFP:BFP screen, including *DGAT2*, *ABHD5*, *ACSL3*, *HILPDA*, *CHP1*, and *AGPAT6* (**Figure 5F,G, Figure S6D,E**), and these genes were absent in the BFP screen (**Figure 5H,I**), indicating that these genes regulate PLIN2 post-translationally. *MARCH6* and *UBE2J2* were also identified as post-translational regulators of PLIN2 (**Figure 5F,G**) in agreement with our earlier studies examining their effects on PLIN2 stability (**Figure 3**). Several interesting novel regulators of PLIN2 were found to act post-translationally (**Figure 5F,G,J,K**), such as the ER scramblase TMEM41B, which was recently implicated in autophagy and LD regulation (Huang et al., 2021; Li et al., 2021; Moretti et al.). Conversely, our screens also identified regulators of PLIN2 expression, which were enriched in genes encoding proteins that localize to the nucleus and function as transcription regulators, such as *HNF1A*, *HNF4A*, *TADA2B*, *NCOA6*, and others (**Figure 5H,I,L,M**). Interestingly, the SREBP pathway components were identified as post-translational regulators of PLIN2, having little effect on PLIN2 transcription (**Figure 5N**). These data argue against PLIN2 as a direct SREBP target and support a model in which SREBP primarily influences PLIN2 stability, likely through its role in controlling cellular lipid metabolism. Some genes that clustered within particular processes, such as mitochondrial fatty acid synthesis, sterol metabolism, oxidative phosphorylation, and protein degradation (**Figure 5O,P; Figure S6F,G**), were detected in both the GFP:BFP and BFP screens, indicating that they have complex roles in regulating both the expression and stability of PLIN2.

To validate these new screens, we further analyzed a small set of candidate regulators. HDAC3 and HNF4A were detected as high confidence regulators based on the enrichment of several sgRNAs (**Figure 6A,E**). Depletion of histone deacetylase 3 (HDAC3) in our PLIN2-GFP-P2A-BFP reporter cells (**Figure S7A**) increased GFP and BFP levels (**Figure 6B**), and parental Huh7 cells depleted of HDAC3 (**Figure S7B**) exhibited increased neutral lipid staining (**Figure 6C**) and increased PLIN2 transcript levels (**Figure 6D**). These data indicate that HDAC3 suppresses PLIN2 expression and are consistent with the known epigenetic regulation of hepatic lipid metabolism by HDAC3 (Sun et al., 2012). Indeed, PLIN2 is upregulated in *HDAC3* null mice, and these mice exhibit hepatic steatosis that can be suppressed by PLIN2 depletion with anti-sense oligos (Sun et al., 2012). Conversely, depletion of Hepatocyte nuclear factor 4-alpha (HNF4A) in PLIN2-GFP-P2A-BFP reporter cells (**Figure S7C**) reduced GFP and BFP (**Figure 6F**), and parental Huh7 cells depleted of HNF4A (**Figure S7D**) exhibited decreased neutral lipid staining (**Figure 6G**) and reduced PLIN2 transcript levels (**Figure 6H**). Confocal imaging further validated the changes in neutral lipids, including an increase in LD size in the HDAC3^KO^ cells and a reduction in LDs in the HNF4A^KO^ cells (**Figure 6I-K**). HNF4A is a transcription factor that regulates hepatocyte differentiation through its roles in controlling the expression of numerous hepatocyte genes associated with glucose and lipid metabolism (Hayhurst et al., 2001; Stoffel and Duncan, 1997). The reduced LDs in the HNF4A^KO^ cells may be due to reduced maintenance of hepatocyte identity. Consistent with this possibility, immunoblotting demonstrates a reduction in the hepatocyte marker FABP1 (**Figure S7D**), and phalloidin fluorescence imaging indicates a remarkable alteration in morphology (**Figure 7L**), with a loss in cobblestone morphology and organized cortical actin staining and an increase in spindle-shaped cells. Moreover, depletion of two additional transcription regulators (*TADA2B* and *SUPT20H*) also led to the predicted reductions in BFP fluorescence, consistent with roles in regulating PLIN2 expression (**Figure S7H-K**. Thus, the screens in the PLIN2-GFP-P2A-BFP reporter cells identify new transcriptional and post-translational PLIN2 regulators.

**Figure 6.**
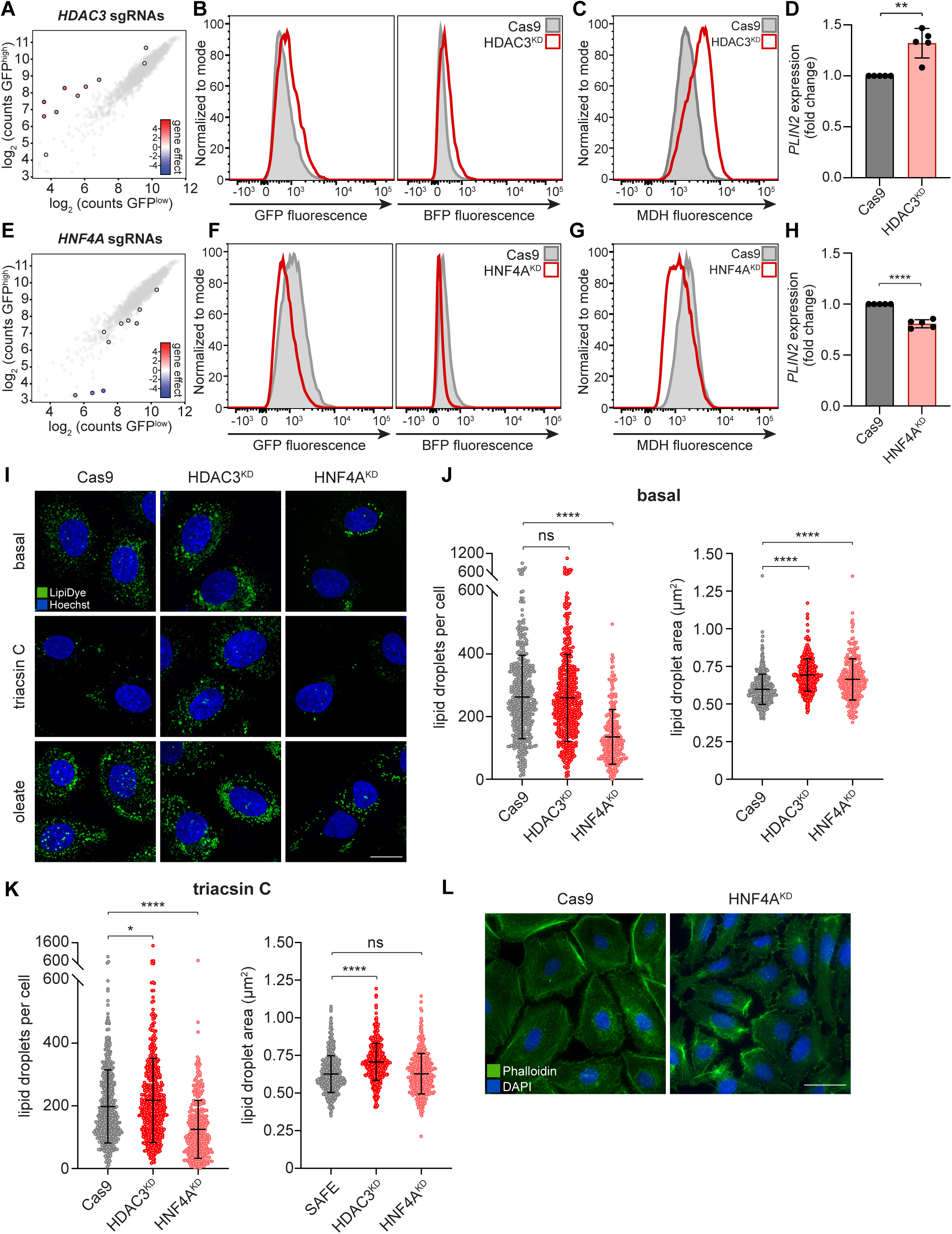
The transcription factor HNF4A regulates expression of PLIN2 and lipid droplet storage. **(A)** Cloud plot indicating deep sequencing counts corresponding to *HDAC3* (color scale) and negative control (grey) sgRNAs from one replicate of the GFP batch retest screen under basal conditions. **(B)** Representative flow cytometry histograms of PLIN2-GFP-P2A-BFP Cas9 cells expressing sgRNAs against a non-coding region of the genome (grey) or *HDAC3* (red). GFP and BFP fluorescence were detected with FITC and Pacific Blue channels, respectively. **(C)** Representative flow cytometry histograms of Huh7 Cas9 cells expressing sgRNAs against a non-coding region of the genome (grey) or *HDAC3* (red). Neutral lipids were stained with 100 µM MDH and fluorescence was measured with the Pacific Blue channel. **(D)** Relative PLIN2 mRNA levels in Huh7 Cas9 cells expressing sgRNAs against a non-coding region of the genome or *HDAC3* as measured by quantitative PCR. Data represent mean ± SD of five biological replicates. ** *p* < 0.01 by two tailed, unpaired t-test. **(E)** Cloud plot indicating deep sequencing counts corresponding to *HNF4A* (color scale) and negative control (grey) sgRNAs from one replicate of the GFP batch retest screen under basal conditions. **(F)** Representative flow cytometry histograms of PLIN2-GFP-P2A-BFP Cas9 cells expressing sgRNAs against a non-coding region of the genome (grey) or *HNF4A* (red). GFP and BFP fluorescence were detected with FITC and Pacific Blue channels, respectively. **(G)** Representative flow cytometry histograms of Huh7 Cas9 cells expressing sgRNAs against a non-coding region of the genome (grey) or *HNF4A* (red). Neutral lipids were stained with 100 µM MDH and fluorescence was measured with the Pacific Blue channel. **(H)** Relative PLIN2 mRNA levels in Huh7 Cas9 cells expressing sgRNAs against a non-coding region of the genome or *HNF4A* as measured by quantitative PCR. Data represent mean ± SD of five biological replicates. **** *p* < 0.0001 by two tailed, unpaired t-test. **(I)** Representative confocal microscopy images of Cas9 control cells expressing sgRNAs against a non-coding region of the genome or sgRNAs against *HDAC3* or *HNF4A*. Cells were imaged under basal conditions or following 8 hr of treatment with 1 µg/ml triacsin C or 200 µM oleate. Lipid droplets were stained with 500 nM Lipi-Green neutral lipid stain and nuclei with 5 µg/mL Hoechst 33342. Scale bar represents 20 µm. **(J)** Quantification of basal lipid droplets per cell (left panel) and LD area (right panel) from (I). Data represent mean ± SD of two biological replicates. \*\*\*\**p* < 0.0001 by one-way ANOVA with Dunnett’s multiple comparisons test. **(K)** Quantification of lipid droplets per cell (left panel) and LD area (right panel) following treatment with 1 µg/ml triacsin C for 8 hr from (I). Data represent mean ± SD of two biological replicates. \**p* < 0.05, \*\*\*\**p* < 0.0001 by one-way ANOVA with Dunnett’s multiple comparisons test. **(L)** Representative fluorescence microscopy image of PLIN2-GFP-P2A-BFP Cas9 cells expressing sgRNAs against a non-coding region of the genome or sgRNAs against *HNF4A*. Actin was stained with Alexa Fluor 488 Phalloidin and nuclei with DAPI. Scale bar represents 50 µm.

**Figure 7.**
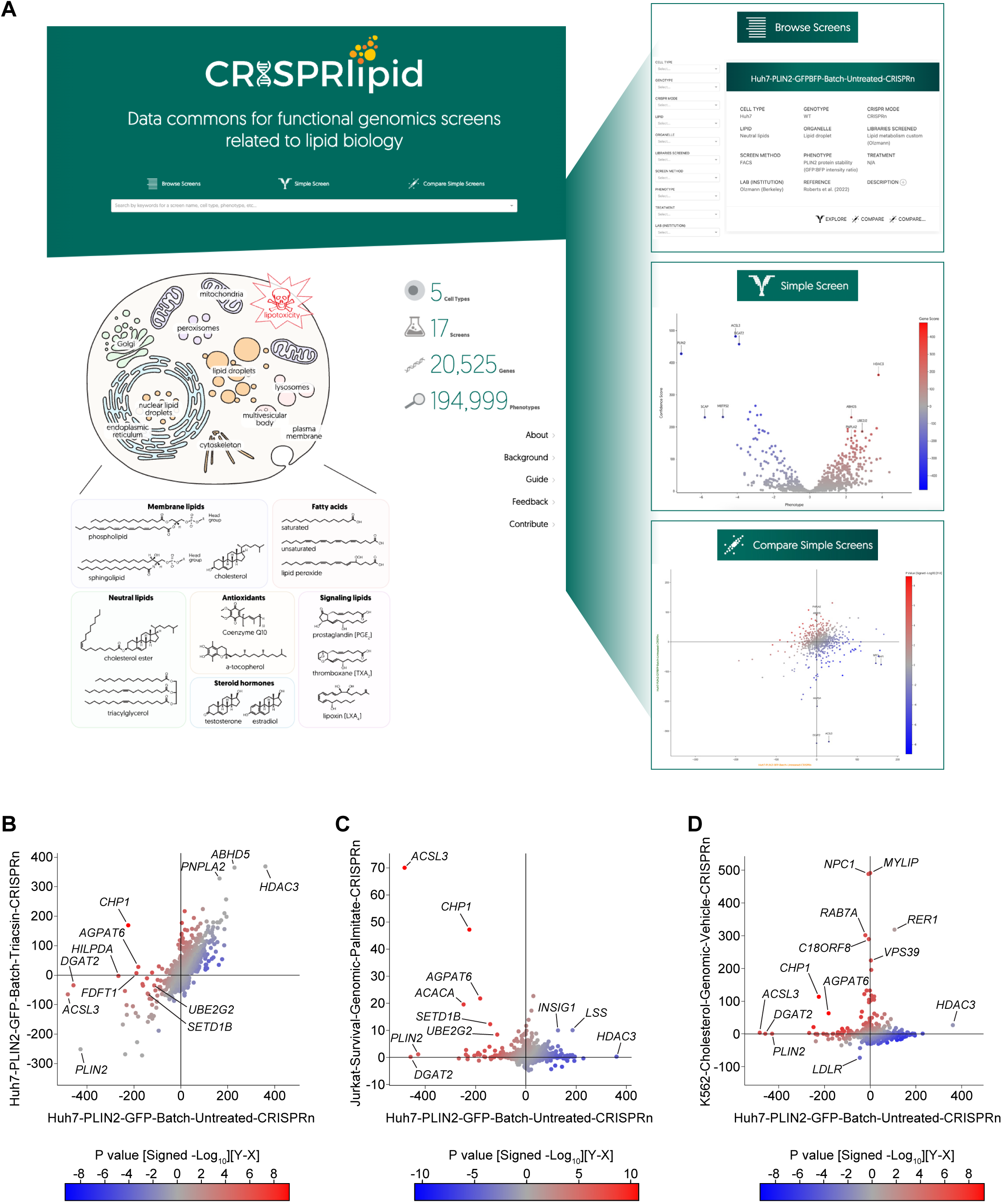
CRISPRlipid: A data commons for functional genomics screens related to lipid biology. **(A)** Landing page for CRISPRlipid (<u>crisprlipid.org</u>), a data repository and tool for sharing, organizing, and analyzing functional genomics screens related to all aspects of lipid biology. Under ‘Browse Screens’ users can explore and search for CRISPR screens based on multiple parameters (e.g., cell type, lipid, organelle, and phenotype). Individual screen datasets (‘Simple Screens’) can be viewed one-by-one and explored with interactive tools, such as marking genes of interest on the plot or selecting subsets of genes that cluster together. Under ‘Compare Simple Screens’, two screens can be plotted on one graph for comparison based on phenotypic or confidence scores. **(B)** Scatter plot of gene scores from FACS-based PLIN2-GFP regulator screen in Huh7 cells under basal conditions (x-axis) versus lipolytic conditions (i.e., triacsin C treated, y-axis). Plot was generated using the CRISPRlipid ‘Compare Simple Screens’ tool. Color scale indicates differential p-value. **(C)** Scatter plot of gene scores from FACS-based PLIN2-GFP regulator screen in Huh7 cells (x-axis) versus palmitate survival screen in Jurkat cells (y-axis). Color scale indicates differential p-value. **(D)** Scatter plot of gene scores from FACS-based PLIN2-GFP regulator screen in Huh7 cells (x-axis) versus FACS-based cholesterol homeostasis screen in K562 cells (y-axis). Color scale indicates differential p-value.

### CRISPRlipid: A data commons for functional genomics screens related to lipid biology

To facilitate the exploration of the functional genomics data generated in this study, we built an online data portal termed CRISPRlipid (http://crisprlipid.org/) (**Figure 7**). This resource provides a community site for data from functional genomics screens related to lipid biology, such as lipid storage and breakdown, lipid signaling, membrane and organelle regulation, and lipotoxicity. CRISPRlipid employs a similar user interface and is conceptually similar to CRISPRbrain (Tian et al., 2021), which focuses on genetic screens in differentiated cell types. Upon its launch, CRISPRlipid contains the data from the PLIN2-GFP and PLIN2-GFP-P2A-BFP screens described in this study, as well as data from published screens that identify genetic modifiers of oxidative lipid damage and ferroptosis (Bersuker et al., 2019; Li et al., 2022), palmitate-induced toxicity (Zhu et al., 2019), and lysosomal cholesterol and bis(monoacylglycerol)phosphate (BMP) levels (Lu et al., 2022). CRISPRlipid allows users to browse screens, enables rapid visualization of screen data as interactive volcano plots and rank plots, and generates pairwise comparisons between different screens (**Figure 7A**). Graphs within CRISPRlipid are interactive, and genes of interest or data from the entire screen can be selected and exported for offline analysis.

One of the most powerful features within CRISPRlipid is the “compare simple screens” tool, which presents pairwise comparisons between different screens as a scatter plot (**Figure 7A**). These comparisons are possible because all screens in CRISPRlipid are analyzed using the same pipeline. For example, we used CRISPRlipid to compare the genetic modifiers of PLIN2-GFP levels under basal conditions with the genetic modifiers of PLIN2-GFP levels under lipolytic conditions (**Figure 7B**), of palmitate toxicity (**Figure 7C**), and of accessible cholesterol (**Figure 7D**). As anticipated, there was a strong correlation between genes that regulate PLIN2-GFP levels under basal and lipolytic conditions, and depletion of genes that regulate LD biogenesis (e.g., *ACSL3* and *DGAT2*) or suppress lipolysis (e.g., *HILPDA*) had strong negative effects on PLIN2-GFP levels under basal conditions but no effect under lipolytic conditions (**Figure 7B**). Interestingly, several factors that positively influence PLIN2-GFP levels and LD biogenesis, ACSL3, CHP1, and AGPAT6, also promote palmitate toxicity (**Figure 7C**). This is unexpected because LDs suppress saturated fatty acid toxicity by mediating their sequestration in TAG (Listenberger et al., 2003). However, ACSL3, CHP1, and AGPAT6 are also involved in early steps of glycerolipid biosynthesis, regulating the biosynthesis of TAG and glycerophospholipids. High amounts of saturated fatty acids in glycerophospholipids reduce membrane fluidity and increase ER stress and cell death. The results in **Figure 7C** indicate that the role of ACSL3, CHP1, and AGPAT6 in incorporating saturated fatty acids into phospholipids is dominant over their protective role in sequestering saturated fatty acids in LDs, consistent with previous findings (Piccolis et al., 2019; Zhu et al., 2019). *ACACA*, *SETD1B*, and *UBE2G2* show a similar functional relationship as *ACSL3*, *CHP1*, and *AGPAT6* (**Figure 7C**), raising the possibility that they play analogous roles in regulating fatty acid flux into glycerolipids. A comparison of the genetic modifiers of PLIN2-GFP levels and accessible cholesterol highlights genes that selectively influence PLIN2-GFP (e.g., *ACSL3*, *DGAT2*) and cholesterol (e.g., *NPC1*, *MYLIP*, *Rab7*, *c18orf8*) along the axes (**Figure 7D**). The comparison also identifies *CHP1* and *AGPAT6* as shared regulators of PLIN2-GFP and cholesterol (**Figure 7D**), raising the possibility that impaired glycerolipid biosynthesis leads to an increase in accessible cholesterol levels. These three pairwise comparisons (**Figure 7B-D**) provide examples of how CRISPRlipid can be used to uncover unexpected functional relationships between genes and to generate new hypotheses of cellular lipid regulation that can be experimentally explored in future studies. In summary, CRISPRlipid provides an extensive community resource for exploration and comparison of functional genomics data related to lipid biology.

## DISCUSSION

In this study, we developed and extensively characterized a set of genome-edited PLIN2 reporter cell lines to study the mechanisms of PLIN2 regulation. Employing these reporter cells, we performed a series of CRISPR-Cas9 loss-of-function screens to globally profile genes that regulate PLIN2 abundance under different metabolic conditions. Following our initial genome-wide screen in the PLIN2-GFP cells, we also generated and screened a new custom LD and Metabolism sgRNA library. Our functional genomics data provide a comprehensive inventory of genes that regulate PLIN2 levels and reveal their mechanism of regulation, transcriptional or post-translational. Given the reciprocal relationship of PLIN2 and LDs, these data also provide a wealth of new regulators of LDs and neutral lipids. We identified not only the expected regulators of LD biogenesis and turnover, but also many new regulators and processes that were not previously connected with PLIN2, such as the electron transport chain and mitochondrial fatty acid synthesis and protein lipolyation pathways. We validated the effects of selected transcription regulators, such as HNF4A and HDAC3, and post-translational regulators, such as MARCH6, UBE2J2, and enzymes within the SREBP pathway, supporting the strength and confidence in our resource.

The identification of MARCH6 and its cognate E2 enzyme UBE2J2 as post-translational regulators of PLIN2 is in line with a previous study that reported MARCH6 ubiquitination and degradation of uninserted PLIN2 (Nguyen et al., 2019). In cell types lacking LDs, MARCH6 recognizes acetylation-dependent degradation signal at the PLIN2 N-terminus, which functions as an N-end rule degron (Nguyen et al., 2019). Although we find that MARCH6 affects PLIN2 levels and LDs in Huh7 cells, our data indicate that MARCH6 indirectly regulates PLIN2 protein stability through its effects on LD abundance. Indeed, loss of MARCH6 still increases LDs in a PLIN2 KO cell line and MARCH6 KO does not affect the rate of PLIN2 clearance during lipolytic breakdown of LDs. The simplest model is that MARCH6 mediates the ubiquitination and degradation of an enzyme involved in neutral lipid synthesis or LD biogenesis. Our data indicate a modest stabilization of SCD1 and ACSL3 (**Figure 4J**), which are known to stimulate LD biogenesis by increasing the amount of activated unsaturated fatty acids, but whether the stabilization of these factors is responsible for the MARCH6 effects on LDs remains to be determined. The question of how LD inserted PLIN2 is degraded during the lipolytic consumption of LDs also remains open. It was recently shown that the E3 ligases UBR1 and UBR2 play redundant roles in promoting PLIN2 proteasomal degradation in hepatocytes (Zhang et al., 2022). It is possible that UBR1 and UBR2 may contribute to PLIN2 degradation under lipolytic conditions. One limitation of CRISPR-Cas9 loss of function screens is the inability to identify genes that can be readily compensated for by redundant pathways. Indeed, TRC8 compensates for the loss of MARCH6 in the degradation of certain substrates (Stefanovic-Barrett et al., 2018). TRC8 was not detected in our screens, but it remains possible that TRC8 or other E3 ligases could partially compensate for the loss of MARCH6. It is notable that induced PLIN2 degradation through overexpression of a constitutively active form of UBR2 was sufficient to suppress hepatic steatosis (Zhang et al., 2022), demonstrating the potential therapeutic value of regulating PLIN2 levels.

With the explosion in “big data”, a major challenge that remains is how to best make data accessible to facilitate discovery. To promote accessibility, exploration, and comparison of data generated from functional genomic screens, we established CRISPRlipid as a new, open access online data commons for the lipid biology community. We envision CRISPRlipid as an open online portal that provides the scientific community with a resource of functional genomics screen data related to diverse aspects of lipid biology, including (but not limited to) neutral lipid storage, lipid biosynthesis and breakdown, lipotoxicity, lipid signaling, lipid enzyme activity, and membrane homeostasis and remodeling. The strength of this site will increase as additional data are deposited from diverse screens in different cell types and under different conditions, and we invite the research community to contribute functional genomics datasets related to lipid biology (broadly defined) to CRISPRlipid. Future iterations of CRISPRlipid will include automated data analysis pipelines and integrated analysis functions across databases that include cell viability data, transcriptomics, and lipidomics. The integration of systems-level datasets provides a new tool for the community and makes an important step towards a comprehensive understanding of the functional networks that govern cellular lipid homeostasis across diverse cell types and conditions.

## ACKNOWLEDGEMENTS

This research was supported by grants from the National Institutes of Health (R01GM112948 and R01DK128099 to J.A.O., F31DK121477 to M.A.R., F32GM113370 to D.E.L., R01GM074874 to R.R.K.) and from the Chan Zuckerberg Initiative Neurodegeneration Challenge Network to M.K. J.A.O. and M.K. are Chan Zuckerberg Biohub Investigators. D.E.L. was also supported by an Alpha-1 Foundation postdoctoral fellowship. F.F. and M.A.N were supported in part by the Intramural Research Programs of the National Institute on Aging (NIA) part of the National Institutes of Health, Department of Health and Human Services; project number Z01-AG000535-014. E.E. was supported by a Cayman Biomedical Research Institute (CABRI) Undergraduate Fellowship.

## AUTHOR CONTRIBUTIONS

M.A.R. and J.A.O. conceived of the project, designed the experiments, wrote the majority of the manuscript. All authors read, edited, and contributed to the manuscript. J.S. and K.K.D. contributed to the generation of reporter cell lines. M.A.R. performed the genetic screens with assistance from D.E.L. and D.W.M. CRISPRlipid resource was built together with M.A.N., F.F., A.J.M., N.T., and M.K. M.L. performed and analyzed the lipidomics experiments. K.H. assisted with FACS and E.E. assisted in the validation of candidate regulators. We are grateful to Drs. Kivanc Birsoy (Rockefeller University), Suzanne Pfeffer (Stanford University) and Albert Lu (University of Barcelona) for contributing their published functional genomics screen data sets for CRISPRlipid. We thank Dr. Zhipeng Li for helpful discussions and critical reading of the manuscript. We thank Doug Guastaferro and Naveed Ziari for technical assistance with casTLE analysis.

## AUTHOR INFORMATION

Correspondence and requests for materials should be addressed to J.A.O.

## METHODS

### Cell Culture and Transfections

Huh7 and HEK293T cells were cultured in DMEM containing 4.5 g/l glucose and l-glutamine (Corning) supplemented with 10% fetal bovine serum (FBS) (Thermo Fisher Scientific and Gemini Bio Products) at 37°C and 5% CO2.

Huh7 siRNA transfections were performed according to manufacturer’s instructions using DharmaFECT 4 Transfection Reagent (Horizon, T-2004-02). Negative control siRNAs were from Qiagen (#1022076), and SQLE siRNAs from Horizon (#L-009646-00-0005, ON-TARGETplus Human SQLE (6713) siRNA - SMARTpool). SQLE SMARTpool sequences are as follows:

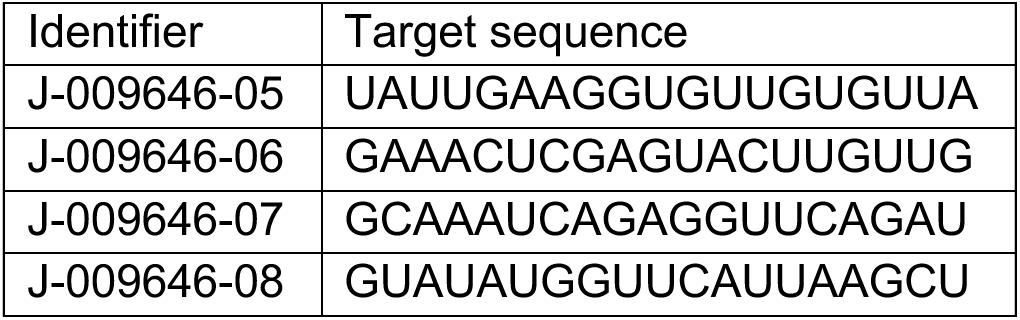

### Generation of GFP-labeled Endogenous PLIN2 Reporter Cells

Endogenous PLIN2 in Huh-7 human liver cells was labeled with a C-terminal AGSGA (flexible linker)-eGFP-S-tag (KETAAAKFERQHMDS) with an internal ribosome entry site (IRES) to allow parallel mTag BFP expression introduced using CRISPR-Cas9 homology-directed repair. The donor construct was produced by first amplifying the target homology region from Huh7 genomic DNA with outer nested primers ns PLIN2 C HDR F 5′-ATGTCCCCAACAACAACTGAGG-3′ and ns PLIN2 C HDR R 5′-CAGATCACTTGAGCCCAGGAAT-3′, followed by phosphorylated inner nested primers PLIN2 HDR F 5′-TTAACTGGTTCCCTGGCAAGAA-3′ & PLIN2 HDR R 5′-CTAGTTGCTCGAGAGGCTGAGG-3′. The phosphorylated fragment was then ligated into a pM575 backbone. A 2062 bp fragment encoding an AGSGA-GFP-S IRES BFP cassette was introduced into the homology region of the intermediate plasmid using polymerase incomplete primer extension cloning with insert primers pI PLIN2 C F 5′- TCTGAGCATAAAACTCATGCCGGCAGCGGCGCCGTGAGCAAG-3′ and pI PLIN2 C R 5′- GTGATAGAGACAGATTTAATTAAGCTTGTGCCCCAGTTTGC-3′ and vector primers pV PLIN2 C F 5′-CACGGCGCCGCTGCCGGCATGAGTTTTATGCTCAGATCGCTG-3′ and pV PLIN2 C R 5′-CTGGGGCACAAGCTTAATTAAATCTGTCTCTATCACTAGTGCATGCTGTGG-3′ to yield the complete pPLIN2 GFP S IRES BFP HDR donor construct, which has 1045 bp and 1231 bp genomic DNA homology arms on either side of the 2062 bp labelling cassette.

CRISPR guide oligos PLIN2 C g1 F 5′-CACCGCATGCACTAGTGATAGGGGC-3′ and PLIN2 C g1 R 5′-AAACGCCCCTATCACTAGTGCATGC-3′ were annealed and ligated into a px330 backbone to yield px330-PLIN2 C g1.

Huh-7 cells were transfected with with 2.67 µg of px330-PLIN2 C CRISPR guide 1 and 5.33 µg of pPLIN2 GFP S IRES BFP HDR donor construct with 8 µL of XtremeGENE HP and treated with 1 uM SCR7 (DNA Ligase IV inhibitor that blocks nonhomologous end-joining) after 6 hr. After 72 hr the media was replenished supplemented with 1 uM SCR7. After a further five days the population was enriched for the 0.05% of cells exhibiting green fluorescence using fluorescence-activated cell sorting (FACS). Cells were sorted another three times to enrich for GFP^+^ cells and eliminate non-GFP^+^ wild type cells from the population.

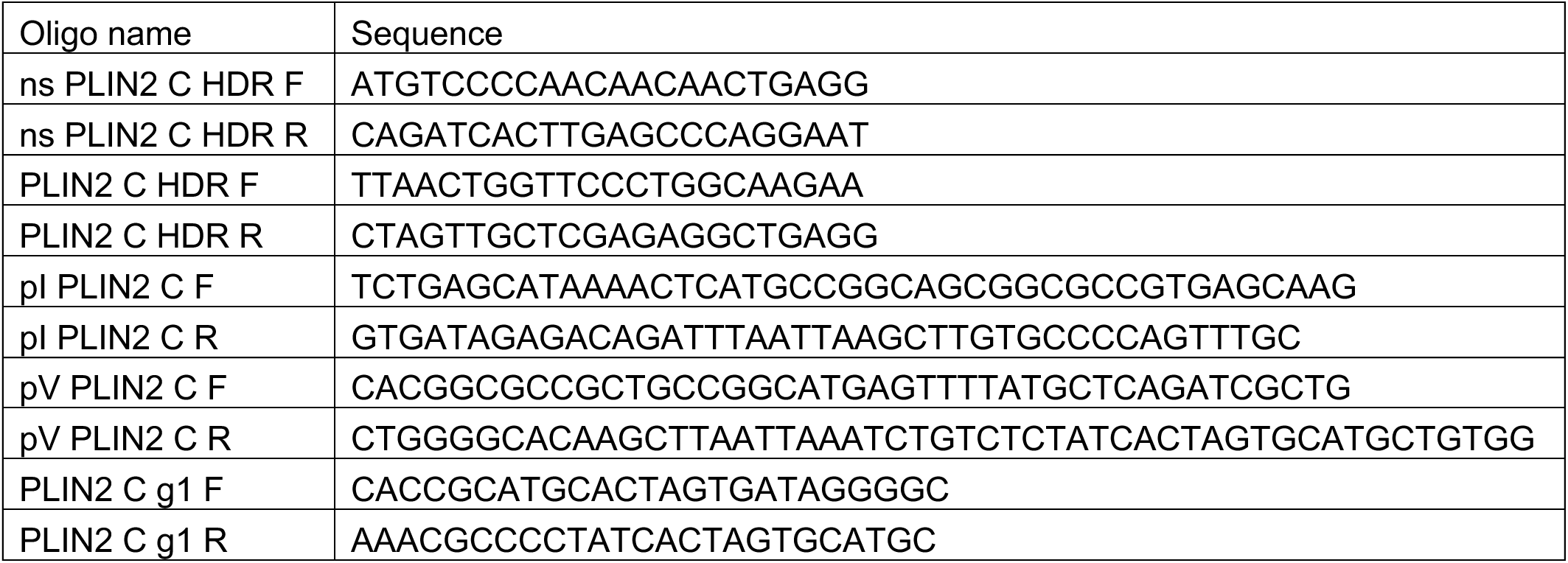

### Generation of GFP-P2A-BFP Endogenous PLIN2 Reporter Cells

To generate the PLIN2-GFP-P2A-BFP knock-in donor plasmid, 500-bp homology arms flanking the PLIN2 stop codon were amplified from Huh7 genomic DNA and inserted in pUC19. 15-bp overlap with each of the PLIN2 homology arms were introduced to the codon optimized GFP-P2A-BFP insert (gBlock synthetic DNA, Integrated DNA Technologies). This insert was then cloned in frame with the PLIN2 stop codon using restriction enzyme-independent fragment insertion by megaprimer cloning. The protospacer adjacent motif site that corresponds to *PLIN2* sgRNA was subsequently mutated in the donor sequence using site-directed mutagenesis primers to prevent cutting of the integrated donor sequence by Cas9.

CRISPR single-guide RNA (sgRNA) sequences targeting *PLIN2* were designed using the CRISPR guide design tool by Benchling (https://www.benchling.com). Underlined nucleotides show the overhangs introduced into oligonucleotides that are necessary for cloning into the BbsI restriction site of vector px330. (Addgene #42230).

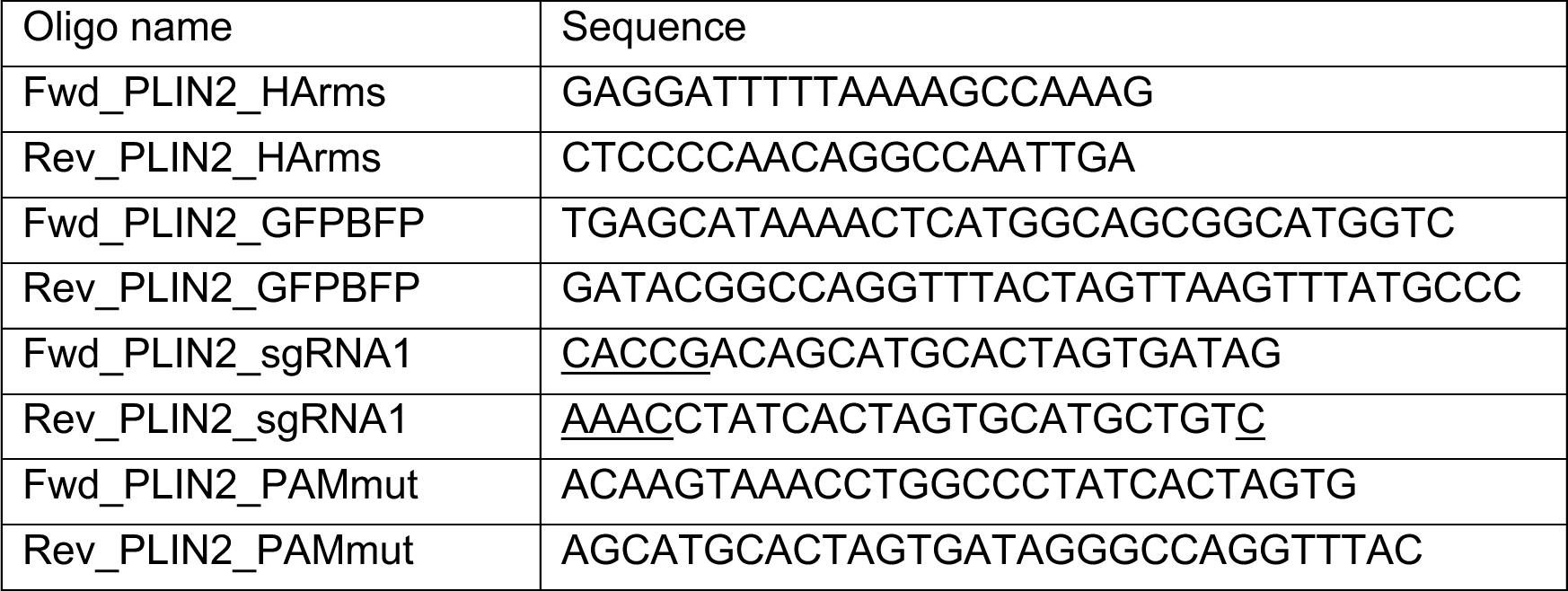

PLIN2 GFP-P2A-BFP knock-in cell lines were generated by co-transfection of Huh7 cells with donor plasmid pUC19 and px330 encoding *PLIN2* sgRNA guide 1 at a 3:1 w/w ratio using X-tremeGENE HP. 6hr post-transfections. Cells were treated and maintained in 1 μM SCR7 for one week, followed by sequential FACS enrichments of GFP^+^/BFP^+^ cells.

### Plasmids

All knockout cell lines were generated using the pMCB320 plasmid, a gift from M. Bassik (Addgene #89359). Guide sequences were selected from the Bassik Human CRISPR Knockout Library (#101926, #101927, #101928, #101929, #101930, #101931, #101932, #101933, #101934) based on enrichment and significance in CRISPR-Cas9 screens. Guide sequences were cloned into pMCB320 using the restriction enzymes BstXI and BlpI.

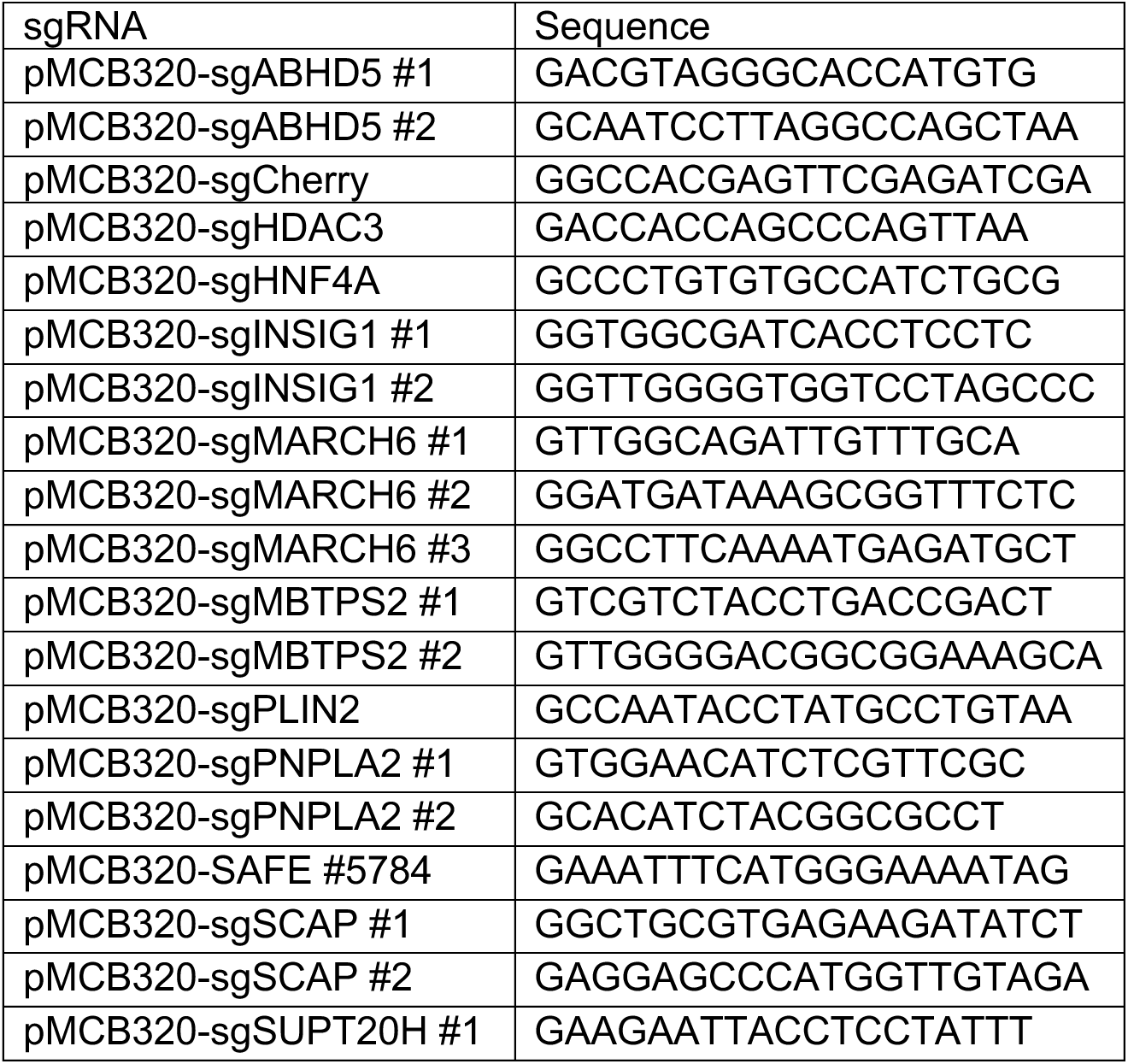
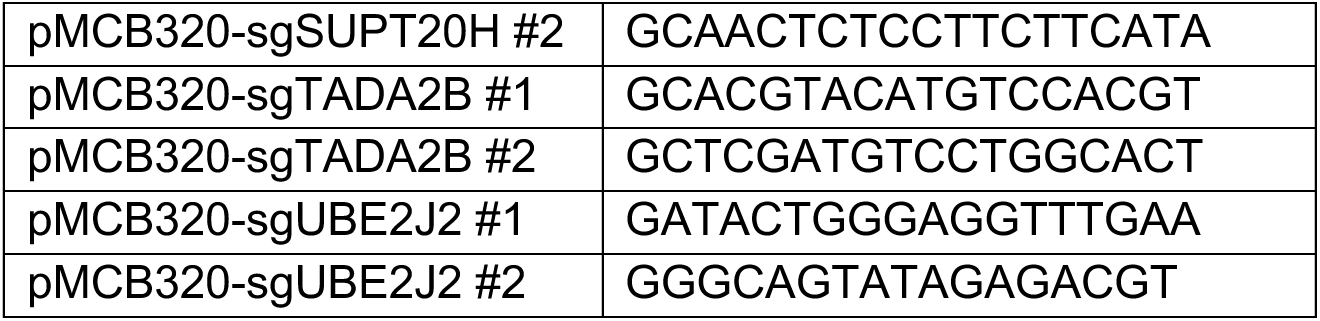

### Generation of CRISPR-Cas9 Genome Edited Cell Lines

For CRISPR-Cas9 screens and individual knockout cell lines, Huh7 wild type, PLIN2-GFP, or PLIN2-GFP-P2A-BFP lines stably expressing Cas9 were generated by infection with lentiCas9- Blast, a gift from F. Zhang (Addgene #52962), and cells were selected in medium containing 4 ug/ml blasticidin. Active Cas9 expression was validated by flow cytometry analysis following infection a self-cleaving mCherry plasmid (pMCB320 which expresses mCherry and an sgRNA targeting the mCherry gene).

Individual knockout lines were generated in Huh7 wild type (HDAC3^KO^, HNF4A^KO^, MARCH6^KO^, PLIN2^KO^,), PLIN2-GFP (ABHD5^KO^, INSIG1^KO^, MARCH6^KO^, MBTPS2^KO^, PNPLA2^KO^, SCAP^KO^, UBE2J2^KO^), and PLIN2-GFP-P2A-BFP (HDAC3^KO^, HNF4A^KO^, SUPT20H^KO^, TADA2B^KO^) cells. To generate lentiviral particles, sgRNA-containing pMCB320 plasmids were cotransfected with third-generation lentiviral packaging plasmids (pVSVG, pRSV and pMDL) into HEK293T cells. Media containing lentivirus was collected 48 or 72 hr after transfection, filtered, and then used to infect cells stably expressing Cas9. After 72 hr of growth, infected cells were selected in media containing 2 μg/ml puromycin (or 0.25 μg/ml hygromycin B for PLIN2^KO^ cells) until over 90% cells were mCherry positive. MARCH6^KO^ (both in Huh7 wild type and in Huh7 PLIN2-GFP lines) and PLIN2^KO^ clones were isolated using serial dilutions. Knockout efficiencies were confirmed via immunoblotting and/or sequencing.

### Immunoblotting

Prior to harvest, cells were washed 2x in DPBS and lysed in 1% SDS. Protein concentrations were determined and normalized using the bicinchoninic acid (BCA) protein assay (Thermo Fisher Scientific). Equal amounts of protein by weight were combined with Laemmli buffer, boiled for 5 min at 95 °C, separated on 4–20% polyacrylamide gradient gels (Bio-Rad Laboratories) and transferred onto nitrocellulose membranes (Bio-Rad Laboratories). Membranes were incubated in 5% nonfat milk in PBS with 0.1% Tween-20 (PBST) for 30 min to reduce nonspecific antibody binding. Membranes were then incubated for at least 2 hr in PBST containing antibodies diluted in 1% BSA, followed by incubation for at least 1 hr in fluorescence-conjugated secondary antibodies diluted in PBST containing 5% nonfat milk. Immunoblots were visualized on a LI-COR imager (LI-COR Biosciences), and ImageJ was used for quantification of protein levels.

### Sequencing of CRISPR-Cas9 Knockouts

Cells were washed 2x and scraped off plates in ice-cold DPBS. Cells were pelleted at 500x g for 5 min. Genomic DNA was extacted and purified using the Qiagen Blood Mini Kit (Qiagen, 51104) according to manufacturer’s instructions. PCR primers were designed using the IDT PrimerQuest Tool (Integrated DNA Technologies). Briefly, primers were designed to flank the predicted Cas9 cut site, with the forward primer ∼100bp upstream of the cut site. PCR was performed on an Applied Biosystems Thermal Cycler using Q5 High-Fidelity 2X Master Mix (New England Biolabs, M0492S). PCR conditions were as follows: 1x 98 °C (2 min), 30x 98 °C (30 sec), 57.5 °C (30 sec), 72 °C (60 sec), 1x 72°C (3 min).

Amplicons were separated on 2% agarose-TAE gels and purified using the QIAquick Gel Extraction Kit (Qiagen). Amplicons were sequenced at Quintarabio, and sequences were assessed for indels using Tracking of Indels by DEcomposition (TIDE, https://tide.nki.nl/) or Inference of CRISPR Edits (ICE; Synthego, https://ice.synthego.com/<u>#/</u>).

### Fluorescence Microscopy

For widefield microscopy, Huh7 PLIN2-GFP cells were grown in 4-well Lab-Tek II Chambered Coverglass (Thermo Fisher Scientific). For live cell imaging of PLIN2-GFP cells, cells were incubated in the presence of 200 µM oleate-BSA complex for 24 hr. Lipid droplets were stained with 0.5 µM Lipi-Deep Red (Dojindo Molecular Technologies) for 2 hours and nuclei were stained with 5 µg/mL Hoeschst 33342 for 30 minutes. Prior to imaging, cells were washed 2x with DPBS and imaged in fresh medium supplemented with 10% FBS lacking phenol red. Live cells were imaged using a Zeiss Axio Observer 7 fitted with a 63x oil objective using DAPI, GFP, and Cy-7 filters. Cells were imaged at 37 °C with 5% CO2. Z-stacks of 0.5-μm thickness were acquired.

For fixed cell imaging of HNF4A^KD^ cells, cells were grown in 12-well plates on glass coverslips coated with poly-l-lysine. Cells were washed 3x with DPBS, fixed for 15 min in DPBS containing 4% (w/v) paraformaldehyde and washed 3x again with DPBS. Cells were permeabilized for 15 min with 1% BSA in DPBS containing 0.1% Triton X-100 and washed 3x with DPBS. Cells were incubated in staining buffer containing Alexa Fluor 488 Phalloidin (Thermo Fisher Scientific, #A12379) in 1% BSA in DPBS for.1 hr in the dark. Cells were washed and then incubated in staining buffer containing DAPI for 20 min in the dark. Cells were washed 3x with DPBS and coverslips mounted on 1 mm glass slides using Fluoromount-G (SouthernBiotech, #0100-01). Cells were imaged using a Zeiss Axio Observer 7 fitted with a 40x objective using DAPI and Alexa Fluor 488 filters. Z-stacks of 0.3-μm thickness were acquired.

For live cell confocal microscopy, cells were grown in 24-well glass bottom plates (170 µm coverglass bottom; Eppendorf, Cellvis). Cells were either left untreated or incubated in the presence of 200 µM oleate-BSA complex or 1 µg/ml triacsin C (Enzo) for 8 hours. Lipid droplets were stained with 0.5 µM Lipi-Green (Dojindo Molecular Technologies) for 2 hours and nuclei were stained with 5 µg/mL Hoeschst 33342 for 30 minutes. Prior to imaging, cells were washed 2x with DPBS and imaged in fresh medium supplemented with 10% FBS lacking phenol red. Live cells were imaged using an Opera Phenix Plus High-Content Screening System (Perkin Elmer) confocal microscope equipped with a 63X water immersion objective using DAPI and GFP filters. Cells were imaged at 37 °C with 5% CO2. Z-stacks of 0.5-μm slices were acquired.

Images were merged and brightness and contrast adjusted using Fiji (https://imagej.net/software/fiji/).

### Lipid Droplet Quantification

LDs were quantified by creating a custom analysis sequence using Harmony High Content Image Analysis Software (Version 4.9). For each field, maximum projection Z-stacks were processed with advanced flatfield correction. Nuclei and cytoplasm were defined using the DAPI and eGFP channels, respectively, and border cells were automatically excluded from analyses. LDs were defined using the “Find Spots” building block (Lipi-Green stain, eGFP channel) thresholding for size, intensity, and roundness. For each cell, lipid droplet number and area (µm2) were quantified. LD quantification data were graphed and analyzed in Prism 9 (GraphPad).

### Genome-wide CRISPR-Cas9 Screens

Genome-wide CRISPR-Cas9 screens were performed using the Bassik Human CRISPR Knockout Library (Morgens et al., 2017). The library consists of nine sublibraries, comprising a total of 225,171 elements, including 212,821 sgRNAs targeting 20,549 genes (∼10 sgRNAs per gene) and 12,350 negative-control sgRNAs. To generate lentiviral particles, each sublibrary was cotransfected with third-generation lentiviral packaging plasmids (pVSVG, pRSV and pMDL) into HEK293T cells. Media containing lentivirus was collected 48 and 72 hr after transfection, combined, and filtered. Huh7 PLIN2-GFP cells stably expressing Cas9 were transduced with lentiviral packaged sublibraries (one sublibrary at a time), with 8 µg/ml polybrene. After 72 hr of growth, infected cells were selected in media containing 2 µg/ml puromycin until over 90% cells were mCherry positive (via flow cytometry). Cells were then recovered for 3-5 days in media lacking puromycin and frozen in liquid nitrogen.

For the screen, library infected cells were thawed (one sublibrary at a time) and expanded at 1,000x coverage (1,000 cells per element). For the steady state genome-wide screen, cells were seeded into 500-cm2 plates (about 10^6 cells per plate, total number of cells seeded was at least 1,000-fold library coverage). For the lipolysis screen, cells were seeded into 500-cm2 plates (about 5 x 10^6 cells per plate). The next day, cells were treated with 1 µg/ml triacsin C for 24 hr. For both the steady state and lipolysis screens, on the day of the sort, cells were dissociated using 0.25% Trypsin-EDTA (Gibco), collected by centrifugation at 300x g for 3 min, and washed 1x with DPBS. Cells were resuspended in phenol red free media (HyClone, 16777-406) supplemented with 3% FBS and 1% BSA (fatty acid free) and kept on ice until FACS.

For both screens, cells were sorted on a BD Aria Fusion equipped with 4 Lasers (488, 405, 561, and 640). The brightest 30% GFP^+^ and dimmest 30% GFP^+^, mCherry^+^ populations were sorted into 15 ml conicals containing DMEM with 4.5 g/l glucose and l-glutamine (Corning) supplemented with 10% FBS. For each sublibrary sort, at least 1000x as many cells as guides were collected. Sorted cells were collected by centrifugation at 1,000g for 10 min, washed 1x with DPBS, and pellets frozen at −80°C until genomic DNA extractions.

Genomic DNA was extracted using the QIAamp DNA Blood Midi Kit (Qiagen) according to the manufacturer’s instructions, with the following modifications. Guide sequence libraries were prepared from genomic DNA by two rounds of PCR using the Herculase II Fusion DNA Polymerase (Agilent). First, guide sequences were amplified from genomic DNA in the following reaction (per 100 μl reaction): 10 μg genomic DNA, 5x Herculase buffer (20 μl), 100 μM oMCB_1562 (1 μl), 100 μM oMCB_1563 (1 μl), 100 mM dNTPs (1 μl), Herculase II Fusion DNA Polymerase (2 μl), and nuclease-free water (to 100 μl). PCR conditions were as follows: 1x 98°C (2 min), 18x 98°C (30 sec), 59.1°C (30 sec), 72°C (45 sec), 1x 72°C (3 min). Amplicons were indexed using Illumina TruSeq LT adapter sequences (for downstream deep sequencing analysis) in the following reaction (per 100 μl reaction): 5 μl PCR1 reaction, 5x Herculase buffer (20 μl), 100 μM oMCB_1439 (0.8 μl), 100 μM barcoded oMCB_1440 (0.8 μl), 100 mM dNTPs (2 μl), Herculase II Fusion DNA Polymerase (2 μl), and nuclease-free water (to 69.4 μl). PCR conditions were as follows: 1x 98°C (2 min), 20x 98°C (30 sec), 59.1°C (30 sec), 72°C (45 sec), 1x 72°C (3 min).

PCR products were separated on a 2% tris-borate-EDTA (TBE)-agarose gel, purified using the QIAquick Gel Extraction Kit (Qiagen) and assessed for quality using a Fragment Analyzer (Agilent). PCR amplicons from each sublibrary-pair (GFP^high^ and GFP^low^) were pooled at a ratio based on sample concentrations (as determined by Qubit Fluorometric Quantification) and the number of elements in each sublibrary. sgRNA sequences were analyzed by deep sequencing using the standard Illumina indexing primer and the custom sequencing primer oMCB1672 (5’- GCCACTTTTTCAAGTTGATAACGGACTAGCCTTATTTAAACTTGCTATGCTGTTTCCAGCTTAGCTCTTAAAC-3’) on an Illumina NextSeq instrument at the Oklahoma Medical Research Foundation.

Sequence reads were aligned to the sgRNA reference library using Bowtie software. For each gene, a gene effect and score (likely maximum effect size and score) and *p*-values were calculated using the Cas9 high-Throughput maximum Likelihood Estimator (casTLE) statistical framework as previously described (Morgens et al., 2016). Deep sequencing counts from the genome-wide screens in PLIN2-GFP reporter cells are available in supplemental tables (**Tables S3,S4**).

### Custom Lipid Droplet and Metabolism sgRNA library construction

A custom “Lipid Droplet and Metabolism” library was generated, consisting of a total of 13,920 elements, including 11,920 sgRNAs targeting 1,196 genes (∼10 sgRNAs per gene) and 2,000 negative control sgRNAs. Guide sequences were from the Bassik Human CRISPR Knockout Library (Morgens et al., 2017) and library construction protocol was previously described (Morgens et al., 2016, 2017).

Briefly, oligonucleotides were synthesized by Twist Biosciences and consist of the guide sequence flanked by restriction enzyme sites and PCR primer sites. The lyophilized oligo pool was resuspended in Tris-HCl (pH 8) and PCR-amplified using KAPA HiFi HotStart DNA Polymerase (Roche, KK2502) with 52 °C anneal and 15 sec extension (10 cycles). PCR products were pooled and purified over a Qiagen MinElute PCR Purification Kit column (Qiagen, 28004) according to manufacturer’s instructions. Eluted sample was restriction digested with BstXI/BlpI overnight at 37 °C. Insert sample was run on a 20% native PAGE gel (Novex TBE Gels, Thermo Fisher Scientific, EC63155BOX), and the band (33 bp) was excised and purified over a Costar Spin X column (Corning, CLS8160). Insert was isopropanol precipitated and resuspended in Qiagen EB. Library vector (pMCB320, Addgene #89359) was restriction digested with BstXI/BlpI for 1 hr at 37 °C, followed by gel purification. Insert was ligated to digested vector (50 ng digested vector: 1 ng digested insert) for 16 hr at 16 °C.

Ligation products were transformed into Lucigen Endura Electrocompetent Cells (Lucigen, 60242) using a Gene Pulser II (Bio-Rad Laboratories) according to manufacturer’s instructions using the following electroporation conditions: 1.8 kV, 600 Ω, 10 µF. Cells were recovered for 2 hr at 37 °C, plated on 500-cm2 agar plates with 100 µg/ml carbenicillin, and grown overnight at 37 °C. The next day, colonies were scraped off plates and library plasmids purified using a Qiagen HiSpeed Maxi kit (Qiagen, 12662). Plasmids were eluted in Qiagen Buffer TE, aliquoted, and stored at −80°C.

For deep sequencing, library plasmids were amplified and indexed using Illumina TruSeq LT adapter sequences in the following reaction (per 100 μl reaction): ∼10 ng purified library plasmid pool, 5x Herculase buffer (20 μl), 100 μM oMCB_1439 (0.8 μl), 100 μM barcoded oMCB_1440 (0.8 μl), 100 mM dNTPs (2 μl), Herculase II Fusion DNA Polymerase (2 μl), and nuclease-free water (to 100 μl). PCR conditions were as follows: 1x 98°C (2 min), 20x 98°C (30 sec), 59.1°C (30 sec), 72°C (45 sec), 1x 72°C (3 min).

PCR products were separated on a 2% tris-borate-EDTA (TBE)-agarose gel, purified using the QIAquick Gel Extraction Kit (Qiagen) and assessed for quality using a Fragment Analyzer (Agilent). PCR amplicons were sequenced using the standard Illumina indexing primer and the custom sequencing primer oMCB1672 on an Illumina MiSeq instrument at the Oklahoma Medical Research Foundation. Sequence reads were aligned to the custom Lipid Droplet and Metabolism sgRNA reference library using Bowtie software and library element distribution was assessed using the casTLE plotDist.py function (Morgens et al., 2016). A list of genes and guides from the Lipid Droplet and Metabolism library is available in **Table S2**.

### Batch Retest CRISPR-Cas9 Screens

Batch retest CRISPR-Cas9 screens were performed using the custom Lipid Droplet and Metabolism Library (this publication). The library consists of 13,920 elements, including 11,920 sgRNAs targeting 1,196 genes (∼10 sgRNAs per gene) and 2,000 negative-control sgRNAs. To generate lentiviral particles, each sublibrary was cotransfected with third-generation lentiviral packaging plasmids (pVSVG, pRSV and pMDL) into HEK293T cells. Media containing lentivirus was collected 48 and 72 hr after transfection, combined, and filtered. Huh7 PLIN2-GFP or PLIN2-GFP-P2A-BFP cells stably expressing Cas9 were transduced with lentiviral packaged library with 8 µg/ml polybrene. After 72 hr of growth, infected cells were selected in media containing 2 µg/ml puromycin until over 90% cells were mCherry positive (via flow cytometry). Cells were then recovered for 3-5 days in media lacking puromycin and frozen in liquid nitrogen.

For each screen, cells were thawed and expanded at >1,000x coverage. For steady state screens, cells were seeded into 500-cm2 plates (about 10^6 cells per plate, total number of cells seeded was roughly 1,000-fold library coverage). For lipolysis screens, cells were seeded into 500-cm2 plates (about 5 x 10^6 cells per plate). The next day, cells were treated with 1 µg/ml triacsin C for 24 hr. For both screens, on the day of the sort, cells were dissociated using 0.25% Trypsin-EDTA (Gibco), collected by centrifugation at 300x g for 3 min, and washed 1x with DPBS. Cells were resuspended in phenol red free media (HyClone) supplemented with 3% FBS and 1% BSA (fatty acid free) and kept on ice until FACS.

For both screens, cells were sorted on a BD Aria Fusion equipped with 4 Lasers (488, 405, 561, and 640). mCherry^+^ cells were sorted as follows: GFP screens (brightest 30% GFP^+^ and dimmest 30% GFP^+^), BFP screens (brightest 30% BFP^+^ and dimmest 30% BFP^+^), GFP:BFP screens (brightest 30% GFP:BFP ratio and dimmest 30% GFP:BFP ratio).Cells were sorted into 15 ml conicals containing DMEM with 4.5 g/l glucose and l-glutamine (Corning) supplemented with 10% FBS. For each sort, at least ∼14 x10^6^ cells were collected (1,000x coverage). Sorted cells were collected by centrifugation at 1,000g for 10 min, washed 1x with DPBS, and pellets frozen at −80°C until genomic DNA extractions.

Genomic DNA was extracted, guide sequences PCR amplified and indexed, and guide enrichment analyzed using casTLE as described above (“Genome-wide CRISPR-Cas9 Screens”). Deep sequencing counts from the batch retest screens are available in supplemental tables (**Tables S5-S7**).

### Flow Cytometry

For analysis of PLIN2-GFP levels, cells were seeded in 6-well plates. The next day, cells were treated with 1 µg/ml triacsin C (Enzo), 200 µM oleate-BSA complex, or DMSO for 24 hr. Cells were dissociated from plates using TrypLE Express (Gibco) and resuspended in DMEM containing 10% FBS. Cells were pelleted by centrifugation at 300x g for 3 min, washed 1X in DPBS, and were resuspended in DPBS. For neutral lipid analysis, cells were incubated with 100 µM of the neutral lipid stain MDH (Yang et al., 2012) (Abcepta) for 30 min on ice prior to pelleting at 300x g.

For all flow cytometry assays, fluorescence was analyzed using an LSRFortessa (BD Biosciences). Levels of PLIN2-GFP were analyzed using the FITC channel, and neutral lipids (MDH) were detected using the Pacific Blue channel. FlowJo Software was used to quantify fluorescence and generate representative histograms.

### Quantitative PCR (qPCR)

RNA was extracted from cells using the Monarch Total RNA Miniprep Kit (New England Biolabs) according to manufacturer’s instructions. Synthesis of cDNA from 100 ng input RNA was performed using the iScript cDNA Synthesis Kit (Bio-Rad Laboratories). Primers were pre-designed (PrimeTime qPCR Primers, Integrated DNA Technologies) and used in conjunction with SsoAdvanced Universal SYBR Green Supermix (Bio-Rad). PCR was performed on CFX96 Touch

Real-Time PCR Detection System (Bio-Rad Laboratories) using the following 3-step amplification protocol: 1x 95 °C (30 sec), 40x 95 °C (10 sec), 60 °C (30 sec), Plate Read. Fold change in mRNA levels were determined using the 2-delta cycle threshold method, normalized to RPLP0 mRNA. IDT predesigned PrimeTime primer pair sequences are as follows:

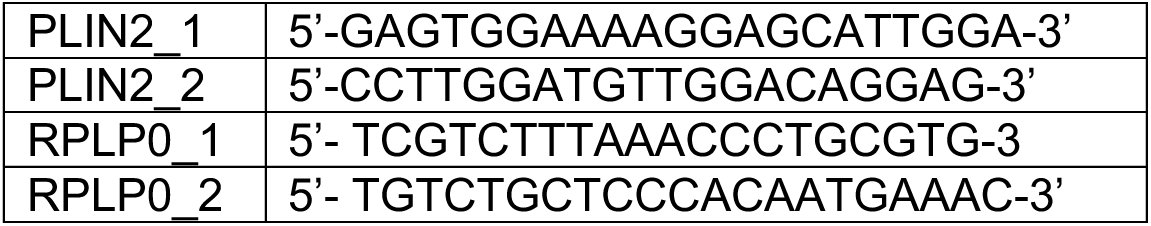

### Lipidomics

#### Chemicals

Materials used for liquid chromatography/mass spectrometry were: water (Optima® LC/MS grade, Fisher Chemical, W6-4), acetonitrile (Optima® LC/MS grade, Fisher Chemical, A955-4), 2-propanol (hypergrade for LC-MS, Supelco, 1.02781.4000), ammonium formate (Lichropur for LC-MS, Sigma-Aldrich, 70221-25G-F) and formic acid (Optima® LC/MS grade, Fisher Chemical, A117-50).

Solvents for lipid extraction were tert-butyl methyl ether (HPLC grade, Sigma-Aldrich, 34875-1L) and methanol (HPLC grade, Fisher Scientific, A452-4) that were spiked with 0.1% (w/v) 2,6-Di-tert-butyl-4-methylphenol (GC grade, Sigma-Aldrich, B1378-100G).

Lipid internal standard mixture was SPLASH ® LIPIDOMIX ® Mass Spec Standard (Avanti Polar Lipids, 330707-1EA).

#### Sample preparation

Approximately 2 million cells were scraped, washed 2x with PBS. Cells were pelleted at 500xg for 5 min, the supernatant was removed, and cell pellets were stored at −80C. Before extraction, cell pellets were thawed on ice for 30 min and resuspended in 50 ul PBS. Internal standards (5 ul SPLASH® LIPIDOMIX® per sample) dissolved in methanol were added directly to each suspension. Lipids were extracted by adding tert-butyl methyl ether (1250 ul) and methanol (375 µL). The mixture was incubated on an orbital mixer for 1 h (room temperature, 32 rpm). To induce phase separation, water (315 µL) was added, and the mixture was incubated on an orbital mixer for 10 min (room temperature, 32 rpm). Samples were centrifuged (room temperature, 10 min, 17,000 x g). Upper organic phase with collected and subsequently dried in vacuo (Eppendorf concentrator 5301, 1 ppm).

#### Liquid Chromatography

Dried lipid extracts were reconstituted in chloroform/methanol (150 ul, 2:1, v/v) and 15 ul of each extract was transferred to HPLC vials containing glass inserts. Quality control samples were generated by mixing equal volumes of each lipid extract followed by aliquotation in 15 ul aliquots. Aliquoted extracts were dried in vacuo (Eppendorf concentrator 5301, 1 ppm) and redissolved in 2-propanol (15 ul) for injection.

Lipids were separated by reversed phase liquid chromatography on a Vanquish Core (Thermo Fisher Scientific, Bremen, Germany) equipped with an Accucore C30 column (150 x 2.1 mm; 2.6 µm, 150 Å, Thermo Fisher Scientific, Bremen, Germany). Lipids were separated by gradient elution with solvent A (acetonitrile/water, 1:1, v/v) and B (2-propanol/acetonitrile/water, 85:10:5, v/v) both containing 5 mM ammonium formate and 0.1% (v/v) formic acid. Separation was performed at 50°C with a flow rate of 0.3 mL/min using the following gradient: 0-15 min – 25 to 86 % B (curve 5), 15-21 min – 86 to 100 % B (curve 5), 21-34.5 min – 100 % B isocratic, 34.5-34.6 min – 100 to 25 % B (curve 5), followed by 8 min re-equilibration at 25 % B.

#### Mass Spectrometry

Reversed phase liquid chromatography was coupled on-line to a Q Exactive Plus Hybrid Quadrupole Orbitrap mass spectrometer (Thermo Fisher Scientific, Bremen, Germany) equipped with a HESI probe. Mass spectra were acquired in positive mode with the following ESI parameters: sheath gas – 40 L/min, auxiliary gas – 10 L/min, sweep gas – 1 L/min, spray voltage – 3.5 kV (positive ion mode); capillary temperature – 250 °C, S-lens RF level – 35 and aux gas heater temperature – 370 °C.

Data acquisition for lipid identification was performed in quality control samples by acquiring data in data dependent acquisition mode (DDA). DDA parameters featured a survey scan resolution of 140,000 (at m/z 200), AGC target 1e6 Maximum injection time 100 ms in a scan range of m/z 240-1200. Data dependent MS/MS scans were acquired with a resolution of 17,500, AGC target 1e5, Maximum injection time 60 ms, loop count 15, isolation window 1.2 m/z and stepped normalized collision energies of 10, 20 and 30 %. A data dependent MS2 was triggered when an AGC target of 2e2 was reached followed by a Dynamic Exclusion for 10 s. All isotopes and charge states > 1 were excluded. All data was acquired in profile mode.

For deep lipidome profiling, iterative exclusion was performed using the IE omics R package (Koelmel et al., 2017). This package generates a list for already fragmented precursors from a prior DDA run that can be excluded from subsequent DDA runs ensuring a higher number of unique MS/MS spectra for deep lipidome profiling. After the initial DDA analysis of a quality control sample, another quality control sample was measured but excluding all previously fragmentated precursor ions. Parameters for generating exclusion lists from previous runs were – RT window = 0.3; noiseCount = 15; MZWindow = 0.02 and MaxRT = 36 min. This workflow was repeated one more time to get a total of three consecutive DDA analyses of a quality control sample in positive ionization mode.

Data for lipid quantification was acquired in Full MS mode with following parameters – scan resolution of 140,000 (at m/z 200), AGC target 1e6 Maximum injection time 100 ms in a scan range of m/z 240-1200.

#### Lipid Identification and Quantification

Lipostar (version 2.1.0, Molecular Discovery, Hertfordshire, UK) equipped with in house generated structure database featuring fatty acids with no information on double bond regio- or stereoisomerism covering glycerolipid, glycerophospholipid, sphingolipid and sterol ester lipid classes. The raw files were imported directly with a Sample MS Signal Filter Signal Threshold = 1000 for MS and a Sample MS/MS Signal Filter Signal Threshold = 10. Automatic peak picking was performed with an m/z tolerance = 5 ppm, chromatography filtering threshold = 0.97, MS filtering threshold = 0.97, Signal filtering threshold = 0. Peaks smoothing was performed using the Savitzky-Golay smoothing algorithm with a window size = 3, degree = 2 and multi-pass iterations = 3. Isotopes were clustered using a m/z tolerance = 5 ppm, RT tolerance = 0.25 min, abundance Dev = 40%, max charge = 1. Peak alignment between samples using an m/z tolerance = 5 ppm and an RT tolerance = 0.25 min. A gap filler with an RT tolerance = 0.05 min and a signal filtering threshold = 0 with an anti Spike filter was applied.

For lipid identification, a “MS/MS only” filter was applied to keep only features with MS/MS spectra for identification. Triacylgylcerols, diacylglycerols and sterol esters were identified as [M+NH4]+ adducts. All phospholipids were identified as [M+H]+ adducts. Following parameters were used for lipid identification: 5 ppm precursor ion mass tolerance and 20 ppm product ion mass tolerance. Automatic approval was performed to keep structures with quality of 3-4 stars. Identifications were refined using manual curation and Kendrick mass defect analysis and lipids that were not following these retention time rules were excluded as false positives.

Quantification by integration of the extracted ion chromatograms of single lipid adducts of these high confidence lipid identifications was manually curated and adjusted. Identified lipids were normalized to peak areas of added internal standards to decrease analytical variation.

For data representation, data was log10 transformed and autoscaled using metaboanalyst.ca (Pang et al., 2022).

## SUPPLEMENTAL TABLES

**Supplementary Table S1. CRISPR/Cas9 Screen Results.**

**Supplementary Table S2. Lipid Droplet and Metabolism Library.**

**Supplementary Table S3. Deep sequencing counts from genome-wide PLIN2-GFP basal screen.**

**Supplementary Table S4. Deep sequencing counts from genome-wide PLIN2-GFP lipolysis screen.**

**Supplementary Table S5. Deep sequencing counts from batch retest PLIN2-GFP basal and lipolytic screens.**

**Supplementary Table S6. Deep sequencing counts from batch retest PLIN2-GFP-P2A-BFP (GFP:BFP sort) basal and lipolytic screens.**

**Supplementary Table S7. Deep sequencing counts from batch retest PLIN2-GFP-P2A-BFP (BFP sort) basal and lipolytic screens.**

---

**Figure S1.**
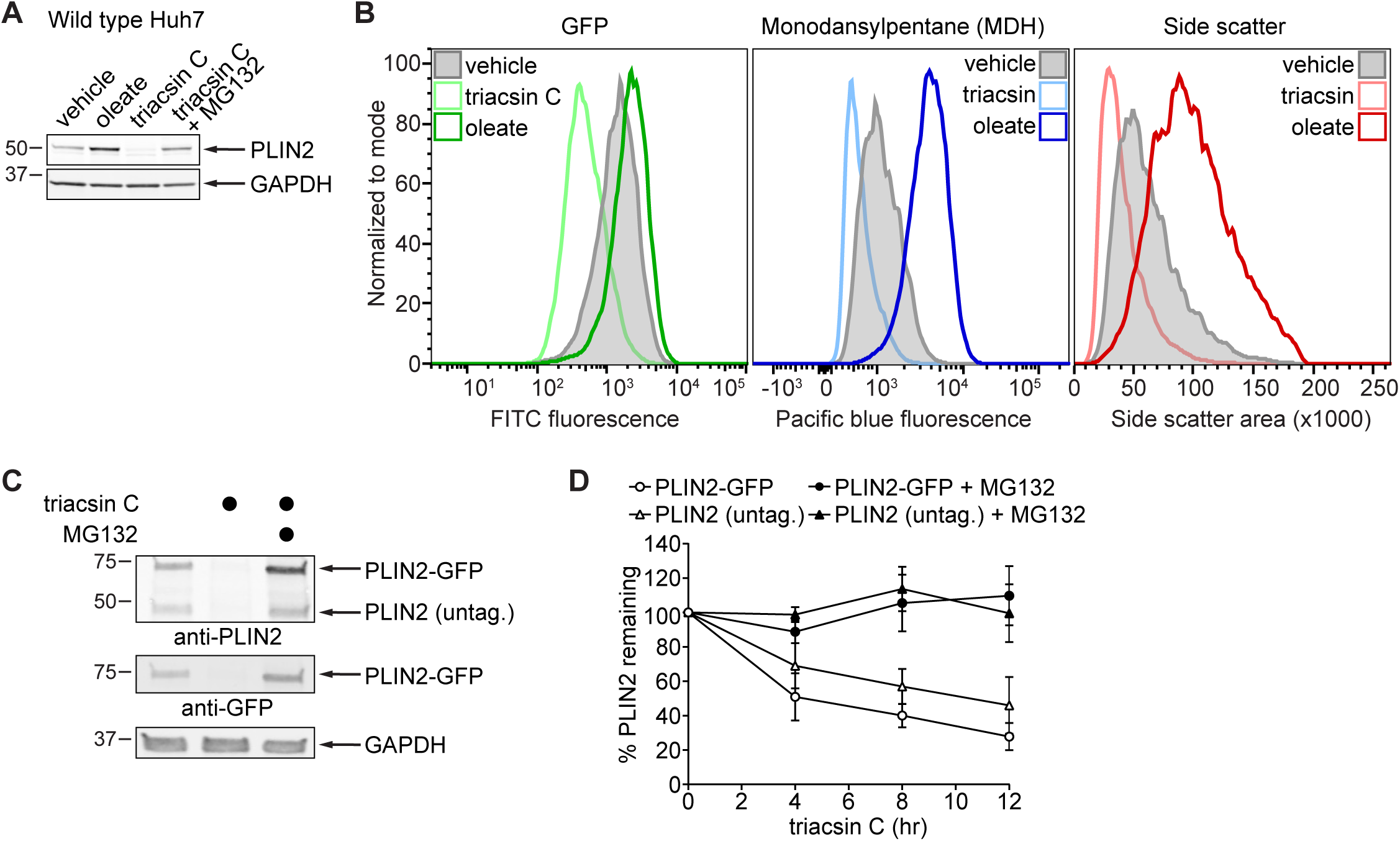
Characterization of PLIN2-GFP reporter cell line. **(A)** Immunoblot of wild type Huh7 cells treated with 200 µM oleate, 1 µg/ml triacsin C, 1 µg/ml triacsin C and 10 µM MG132, or DMSO for 24 hr. Cells were lysed in SDS and immunoblotted for PLIN2 and GAPDH. **(B)** Flow cytometry analysis of PLIN2-GFP cells treated with 1 µg/ml triacsin C or 200 µM oleate for 24 hr and incubated in 100 µM MDH (lipid droplet stain). Levels of PLIN2-GFP and neutral lipids were detected with the FITC and Pacific Blue channels, respectively, and side scatter (area) measured as an indirect readout of lipid droplet abundance. **(C)** Immunoblot of PLIN2-GFP cells treated with 1 µg/ml triacsin C, 1 µg/ml triacsin C with 10 µM MG132, or DMSO for 24 hr. Cells were lysed in SDS and immunoblotted for PLIN2, GFP, and GAPDH. **(D)** PLIN2-GFP cells were treated with 1 µg/ml triacsin, 10 µM MG132, or both for the indicated times. Cells were lysed in SDS and immunoblotted for PLIN2. PLIN2-GFP and PLIN2 protein levels were quantified via densitometry. Data are represented as mean ± SD for three biological replicates.

**Figure S2.**
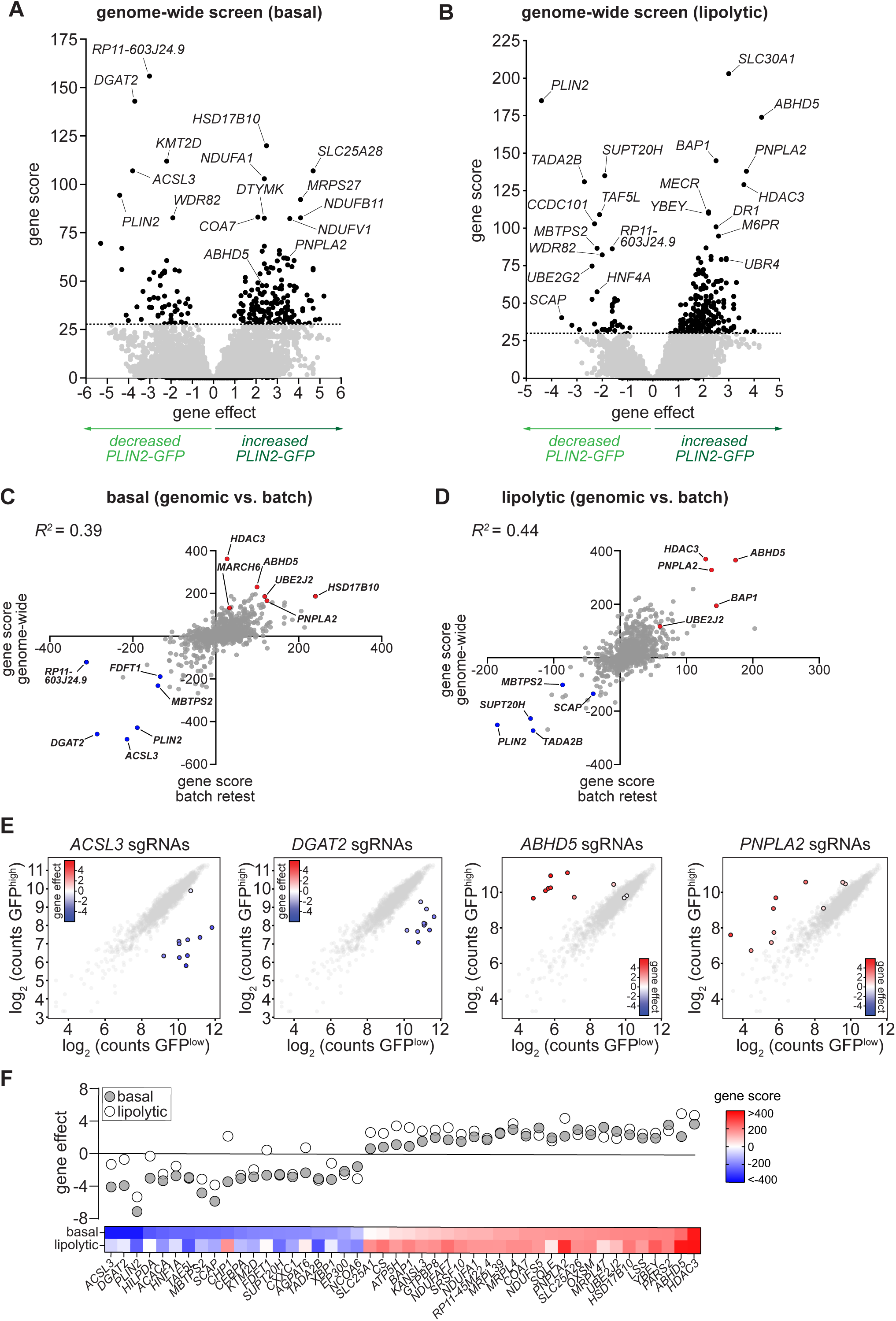
Comparison of genome-wide and batch retest screens in PLIN2-GFP cells. **(A,B)** Volcano plots indicating the gene effects and gene scores for individual genes from genome-wide CRISPR-Cas9 screens of PLIN2-GFP cells under basal (A) and triacsin C treated (B) conditions. Gene effects and gene scores were computed from two biological replicates per screen. Dashed lines indicate the 10 percent false discovery rate cutoff. **(C,D)** Scatter plot of signed gene scores for individual genes from genome-wide screens (x-axis) versus batch retest screens (y-axis) under basal (C) or triacsin C treated (D) conditions. **(E)** Cloud plots indicating deep sequencing counts corresponding to *ACSL3*, *DGAT2*, *ABHD5*, and *PNPLA2* (color scale) or negative control sgRNAs (grey scale) from one replicate of the PLIN2-GFP basal (*ACSL3* and *DGAT2)* or triacsin C treated (*ABHD5* and *PNPLA2*) batch retest screens. **(F)** Heat map and scatter plot displaying the top 30 genes from the basal and triacsin C batch retest screens. Heat map indicates signed gene scores from each screen. Scatter plot indicates gene effects (phenotypic score) of each gene from the basal (grey) and triacsin C (white) screens.

**Figure S3.**
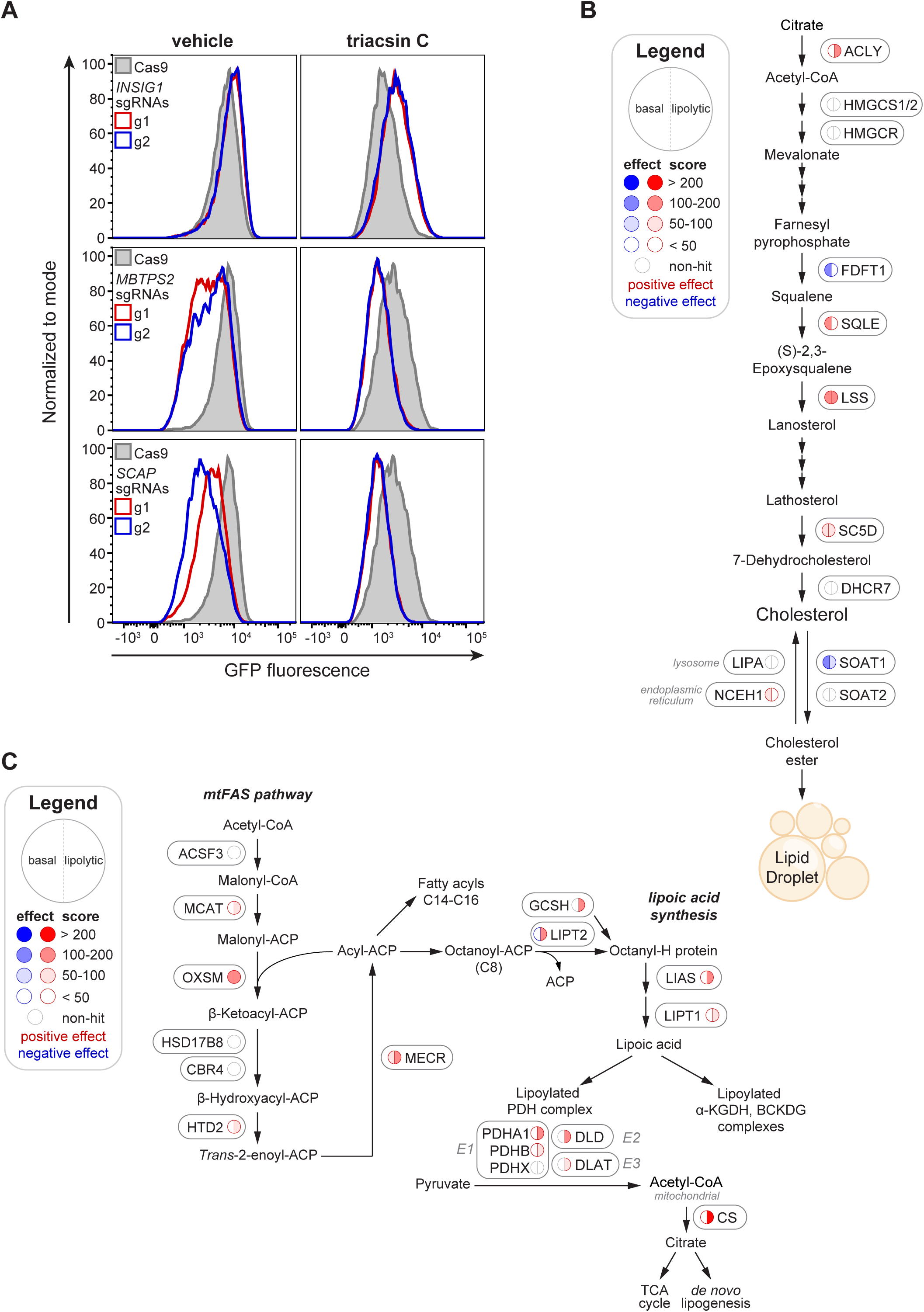
Validation of PLIN2-GFP regulators and biochemical pathway maps. **(A)** Representative flow cytometry histograms of PLIN2-GFP Cas9 cells expressing no sgRNAs (grey) or sgRNAs against *INSIG1* (top), *MBTPS2* (middle), or *SCAP* (bottom). Cells were treated with 1 µg/ml triacsin C (right panels) or DMSO (left panels) for 24 hr. GFP was detected using the FITC channel. **(B)** Schematic of the cholesterol synthesis pathway. Genes are annotated with nodes corresponding to gene effects and scores from batch retest basal and triacsin C screens. **(C)** Schematic of the mitochondrial fatty acid synthesis and lipoic acid synthesis pathways. Genes are annotated with nodes corresponding to gene effects and scores from batch retest basal and triacsin C screens.

**Figure S4.**
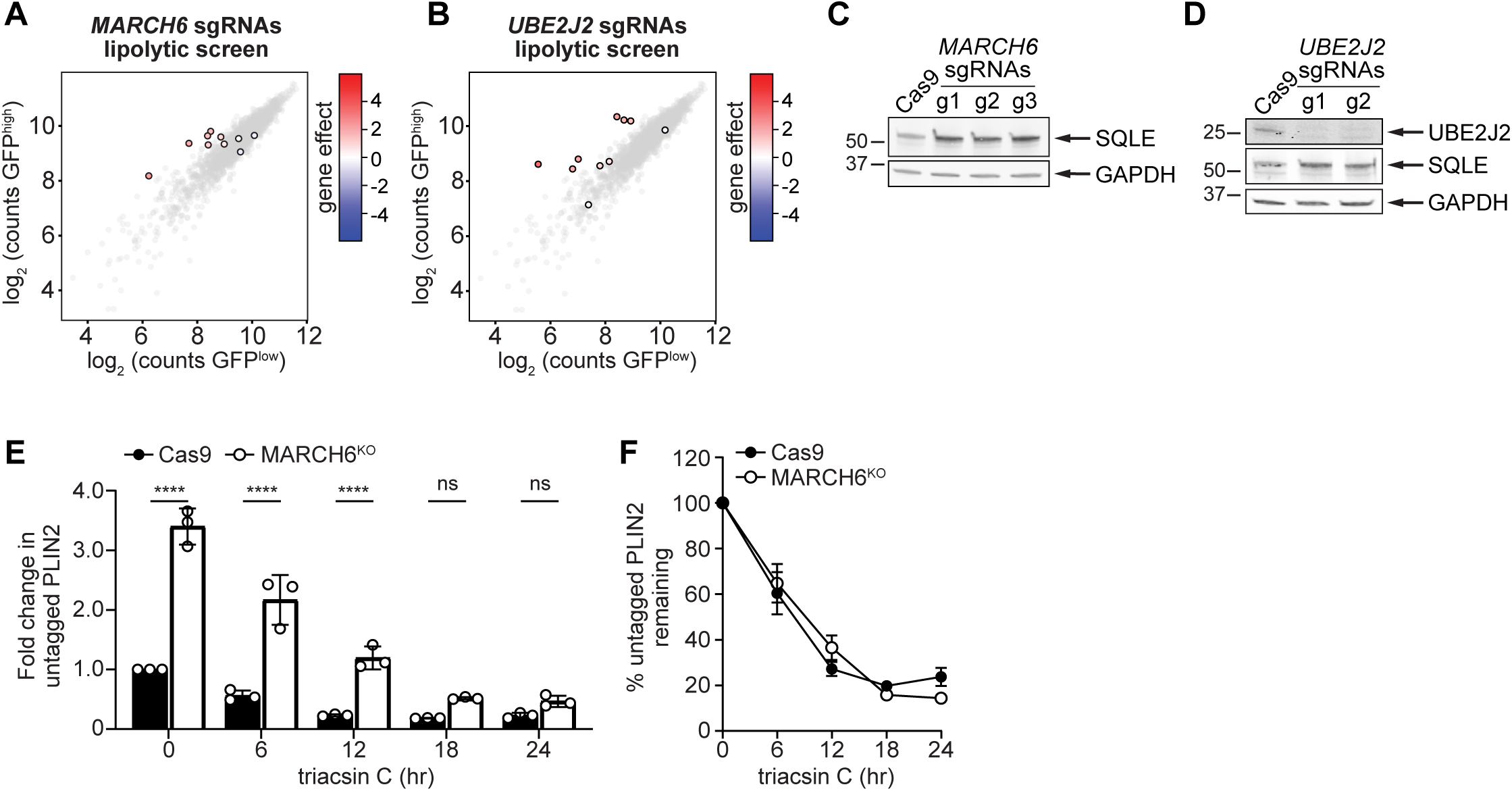
Characterization of MARCH6^KO^ and UBE2J2^KO^ cells. **(A)** Cloud plot indicating deep sequencing counts corresponding to *MARCH6* (color scale) or negative control sgRNAs (grey scale) from one replicate of the PLIN2-GFP batch retest screen under triacsin C treated conditions. **(B)** Cloud plot indicating deep sequencing counts corresponding to *UBE2J2* (color scale) or negative control sgRNAs (grey scale) from one replicate of the PLIN2-GFP batch retest screen under triacsin C treated conditions. **(C)** Immunoblot of Huh7 Cas9 cells expressing no sgRNAs or sgRNAs against *MARCH6* (three different guide pools). Cells were lysed in SDS and immunoblotted for SQLE and GAPDH. **(D)** Immunoblot of Huh7 Cas9 cells expressing no sgRNAs or sgRNAs against *UBE2J2* (two different guide pools). Cells were lysed in SDS and immunoblotted for UBE2J2, SQLE, and GAPDH. **(E)** Quantification of untagged PLIN2 protein levels in PLIN2-GFP Cas9 cells expressing no sgRNAs or sgRNAs against *MARCH6* at the indicated triacsin C treatment times (from Figure 3G). Data represent mean ± SD of three biological replicates. **** p < 0.0001. **(F)** Quantification of untagged PLIN2 protein levels from Figure 3G. PLIN2 protein levels at each time point were normalized to levels at time 0 hr for each cell line. Data represent mean ± SD of three biological replicates.

**Figure S5.**
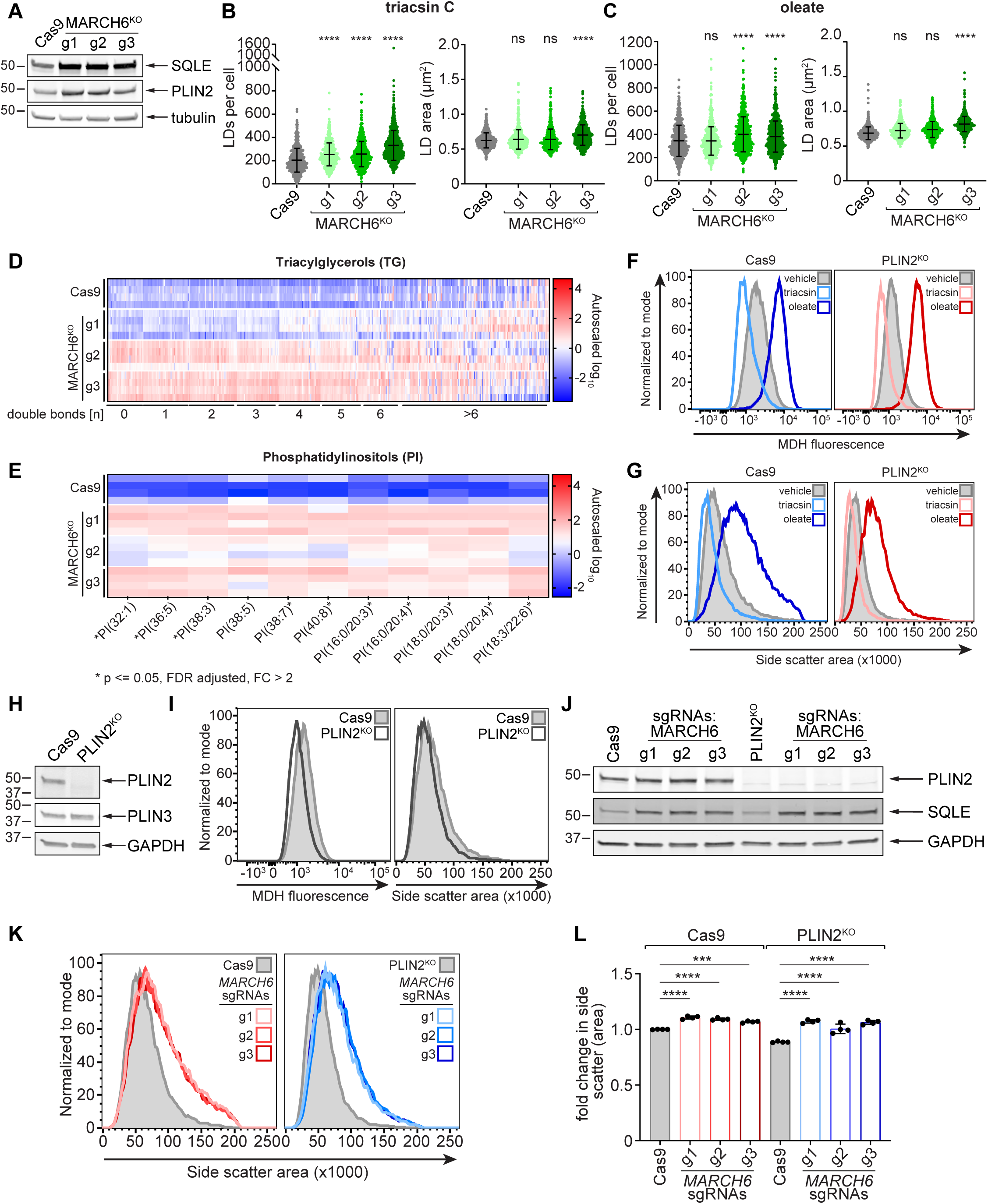
Analysis of the PLIN2 and MARCH6 relationship in LD regulation. **(A)** Immunoblot of Huh7 Cas9 cells expressing sgRNAs against a non-coding region of the genome or sgRNAs against three different sgRNAs against *MARCH6*. Cells were lysed in SDS and proteins separated by SDS-PAGE and immunoblotted for SQLE, PLIN2, and tubulin. **(B)** Quantification of the number (left panel) and area (right panel) of lipid droplets per cell under triacsin C treated conditions from (Figure 4A). Data represent mean ± SD of three biological replicates. \*\*\*\**p* < 0.0001 by one-way ANOVA with Dunnett’s multiple comparisons test. **(C)** Quantification of the number (left panel) and area (right panel) of lipid droplets per cell under oleate treated conditions from (Figure 4A). Data represent mean ± SD of three biological replicates. \*\*\*\**p* < 0.0001 by one-way ANOVA with Dunnett’s multiple comparisons test. **(D)** Heat map indicating changes in triacylglycerol species from the untargeted lipidomics analysis of Cas9 control cells expressing sgRNAs against a non-coding region of the genome and three independent, clonally isolated MARCH6^KO^ cell lines. Data are autoscaled and normalized to the average abundance of each species presented. **(E)** Heat map indicating changes in phosphatidylinositol species from the untargeted lipidomics analysis of Cas9 control cells expressing sgRNAs against a non-coding region of the genome and three independent, clonally isolated MARCH6^KO^ cell lines. Data are autoscaled and normalized to the average abundance of each species presented. **(F)** Flow cytometry histograms of Huh7 Cas9 cells expressing sgRNAs against a non-coding region of the genome or clonally isolated PLIN2^KO^ cells. Cells were incubated in 1 μg/ml triacsin or 200 μM oleate for 24 hr. Neutral lipids were stained with 100 μM MDH and analyzed using the Pacific Blue channel. **(G)** Flow cytometry histograms of Huh7 Cas9 cells expressing sgRNAs against a non-coding region of the genome or clonally isolated PLIN2^KO^ cells. Cells were incubated in 1 μg/ml triacsin or 200 μM oleate for 24 hr. Side scatter (area) was measured as an indirect readout of lipid droplet abundance. **(H)** Immunoblot of Huh7 Cas9 cells expressing sgRNAs against a non-coding region of the genome or clonally isolated PLIN2^KO^ cells. Cells were lysed in SDS and proteins separated by SDS-PAGE and immunoblotted for PLIN2, PLIN3, and GAPDH. **(I)** Flow cytometry histograms of Huh7 Cas9 cells expressing sgRNAs against a non-coding region of the genome or clonally isolated PLIN2^KO^ cells at steady state. Neutral lipids were stained with 100 μM MDH and analyzed using the Pacific Blue channel (left) or side scatter (right). **(J)** Immunoblot of Cas9 control cells expressing sgRNAs against a non-coding region of the genome or three different sgRNAs against *MARCH6*^KO^ (left four lanes) or clonally isolated PLIN2^KO^ cells expressing sgRNAs against a non-coding region of the genome or three different sgRNAs against *MARCH6*^KO^ (right four lanes). Cells were lysed in SDS and immunoblotted for PLIN2, SQLE, and GAPDH. **(K)** Representative flow cytometry histograms measuring side scatter (area) in Huh7 Cas9 cells expressing sgRNAs against a non-coding region of the genome or three different *MARCH6* sgRNAs (left panel) or PLIN2^KO^ cells expressing sgRNAs against a non-coding region of the genome or three different sgRNAs against *MARCH6* (right panel). **(L)** Quantification of the fold change in mean side scatter from (I). Data represent mean ± SD of three biological replicates. *** *p* < 0.001, **** *p* < 0.0001.

**Figure S6.**
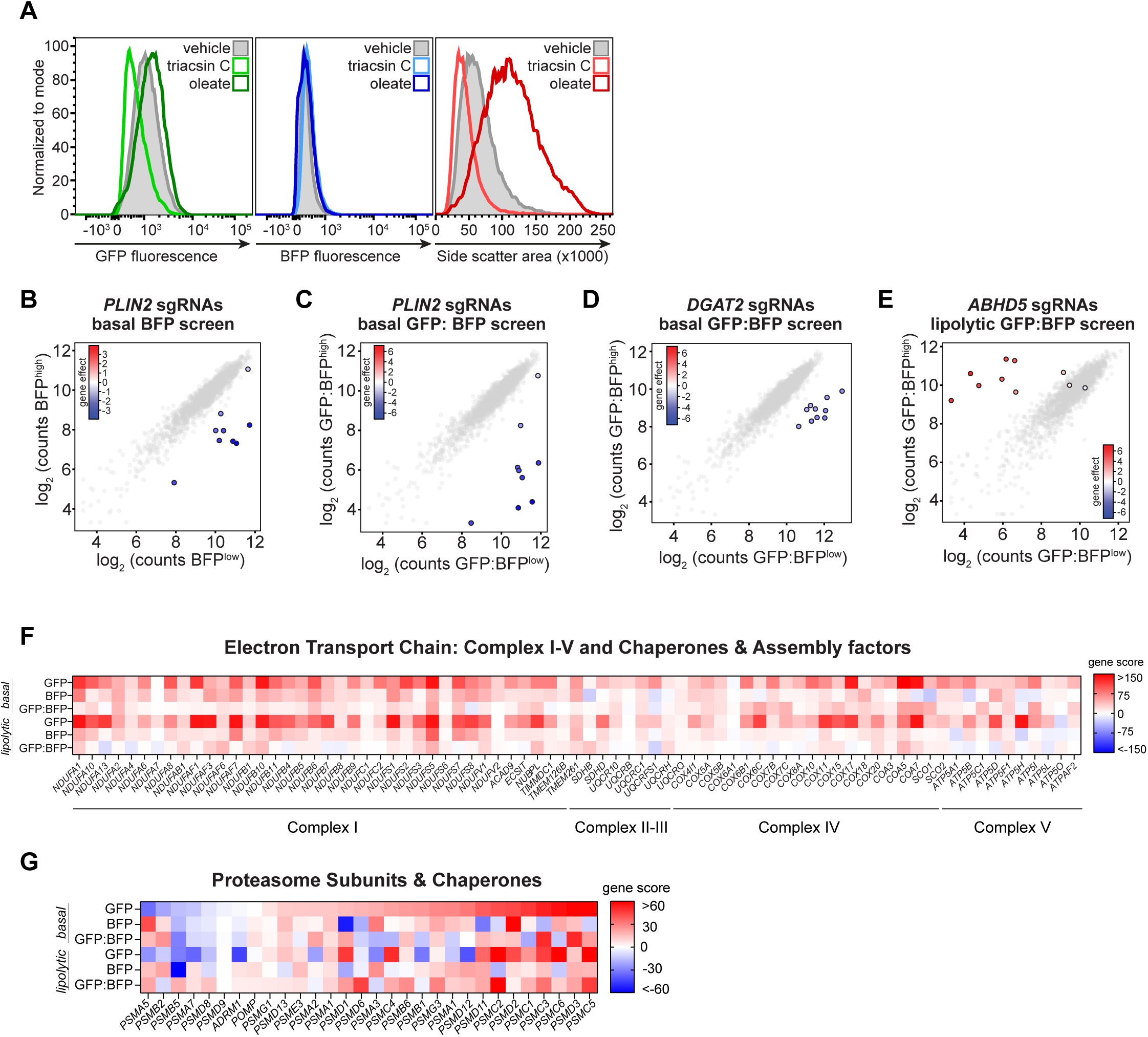
Analysis of PLIN2-GFP-P2A-BFP reporter cells and identified genetic modifiers. **(A)** Flow cytometry analysis of PLIN2-GFP-P2A-BFP cells treated with 1 µg/ml triacsin C or 200 µM oleate for 24 hr and incubated in 100 µM MDH (lipid droplet stain). Levels of PLIN2-GFP and neutral lipids were detected with the FITC and Pacific Blue channels, respectively, and side scatter (area) was measured as an indirect readout of lipid droplet abundance. **(B)** Cloud plot indicating deep sequencing counts corresponding to *PLIN2* (color scale) or negative control sgRNAs (grey scale) from one replicate of the BFP batch retest screen under basal conditions. **(C)** Cloud plot indicating deep sequencing counts corresponding to *PLIN2* (color scale) or negative control sgRNAs (grey scale) from one replicate of the GFP:BFP batch retest screen under basal conditions. **(D)** Cloud plot indicating deep sequencing counts corresponding to *DGAT2* (color scale) or negative control sgRNAs (grey scale) from one replicate of the GFP:BFP batch retest screen under basal conditions. **(E)** Cloud plot indicating deep sequencing counts corresponding to *ABHD5* (color scale) or negative control sgRNAs (grey scale) from one replicate of the GFP:BFP batch retest screen under triacsin C treated conditions. **(F)** Heat map displaying the signed gene scores of electron transport chain-related genes from batch retest GFP, BFP, and GFP:BFP screens. **(G)** Heat map displaying the signed gene scores of proteasome-related genes from batch retest GFP, BFP, and GFP:BFP screens.

**Figure S7.**
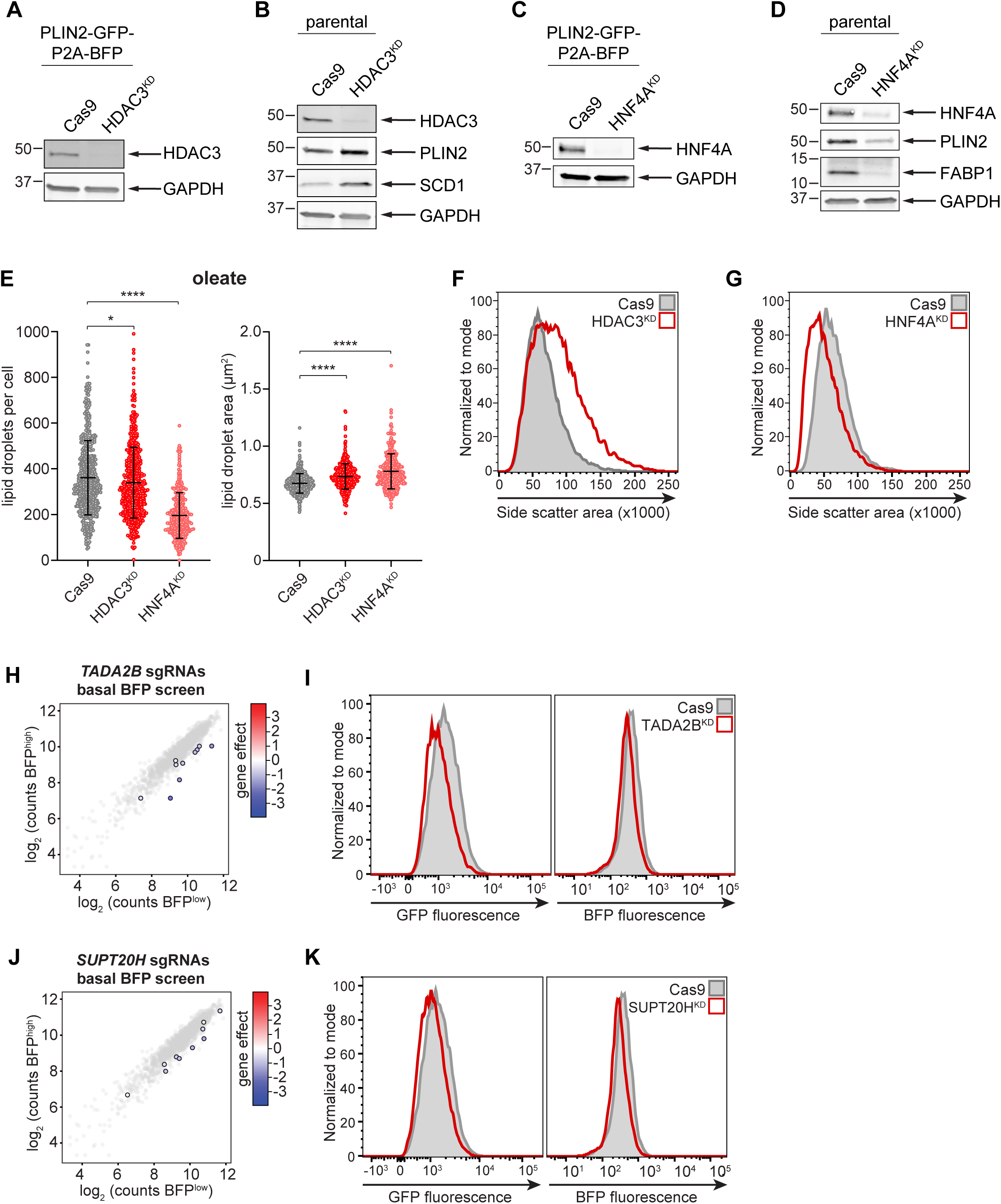
Characterization of genetic modifiers of PLIN2 expression. **(A)** Immunoblot of Huh7 PLIN2-GFP-P2A-BFP Cas9 cells expressing sgRNAs against a non-coding region of the genome or sgRNAs against *HDAC3*. Cells were lysed in SDS and proteins separated by SDS-PAGE and immuoblotted for HDAC3 and GAPDH. **(B)** Immunoblot of parental Huh7 Cas9 cells expressing sgRNAs against a non-coding region of the genome or sgRNAs against *HDAC3*. Cells were lysed in SDS and proteins separated by SDS-PAGE and immunoblotted for HDAC3, PLIN2, SCD1, and GAPDH. **(C)** Immunoblot of Huh7 PLIN2-GFP-P2A-BFP Cas9 cells expressing sgRNAs against a non-coding region of the genome or sgRNAs against *HNF4A*. Cells were lysed in SDS and proteins separated by SDS-PAGE and immunoblotted for HNF4A and GAPDH. **(D)** Immunoblot of parental Huh7 Cas9 cells expressing sgRNAs against a non-coding region of the genome or sgRNAs against *HNF4A*. Cells were lysed in SDS and proteins separated by SDS-PAGE and immunoblotted for HNF4A, PLIN2, FABP1, and GAPDH. **(E)** Quantification of the number (left panel) and area (right panel) of lipid droplets per cell under oleate treated conditions from (Figure 6I). Data represent mean ± SD of two biological replicates. Data represent mean ± SD of two biological replicates. \**p* < 0.05, \*\*\*\**p* < 0.0001 by one-way ANOVA with Dunnett’s multiple comparisons test. **(F)** Flow cytometry histograms of side scatter (area) in Huh7 Cas9 cells expressing sgRNAs against a non-coding region of the genome (grey) or sgRNAs against *HDAC3* (red). **(G)** Flow cytometry histograms of side scatter (area) in Huh7 Cas9 cells expressing sgRNAs against a non-coding region of the genome (grey) or sgRNAs against *HNF4A* (red). **(H)** Cloud plot indicating deep sequencing counts corresponding to *TADA2B* (color scale) and negative control (grey) sgRNAs from one replicate of the BFP batch retest screen under basal conditions. **(I)** Representative flow cytometry histograms of PLIN2-GFP-P2A-BFP Cas9 cells expressing sgRNAs against a non-coding region of the genome (grey) or sgRNAs against *TADA2B* (red). GFP and BFP fluorescence were detected with FITC and Pacific Blue channels, respectively. **(J)** Cloud plot indicating deep sequencing counts corresponding to *SUPT20H* (color scale) and negative control (grey) sgRNAs from one replicate of the BFP batch retest screen under basal conditions. **(K)** Representative flow cytometry histograms of PLIN2-GFP-P2A-BFP Cas9 cells expressing sgRNAs against a non-coding region of the genome (grey) or sgRNAs against *SUPT20H* (red). GFP and BFP fluorescence were detected with FITC and Pacific Blue channels, respectively.

